# The isolate *Caproiciproducens* sp. 7D4C2 produces *n*-caproate at mildly acidic conditions from hexoses: genome and rBOX comparison with related strains and chain-elongating bacteria

**DOI:** 10.1101/2020.07.19.210914

**Authors:** Sofia Esquivel-Elizondo, Caner Bağcı, Monika Temovska, Byoung Seung Jeon, Irina Bessarab, Rohan B. H. Williams, Daniel H. Huson, Largus T. Angenent

## Abstract

**Background:** Bulk production of medium-chain carboxylates (MCCs) with 6-12 carbon atoms is of great interest to biotechnology. Open cultures (*e*.*g*., reactor microbiomes) have been utilized to generate MCCs in bioreactors. When in-line MCC extraction and prevention of product inhibition is required, the bioreactors have been operated at mildly acidic pH (5.0-5.5). However, model chain-elongating bacteria grow optimally at neutral pH values. Here, we isolated a chain-elongating bacterium (strain 7D4C2) that thrives at mildly acidic pH. We studied its metabolism and compared its whole genome and the reverse β-oxidation (rBOX) genes to other bacteria.

**Results:** Strain 7D4C2 produces lactate, acetate, *n*-butyrate, *n*-caproate, biomass, and H_2_/CO_2_ from hexoses. With only fructose as substrate (pH 5.5), the maximum *n*-caproate specificity (*i*.*e*., products *per* other carboxylates produced) was 60.9 ± 1.5%. However, this was considerably higher at 83.1 ± 0.44% when both fructose and *n*-butyrate (electron acceptor) were combined as a substrate. A comparison of serum bottles with fructose and *n*-butyrate with an increasing pH value from 4.5 to 9.0 showed a decreasing *n*-caproate specificity from ∼92% at mildly acidic pH (pH 4.5-5.0) to ∼24% at alkaline pH (pH 9.0). Moreover, when carboxylates were extracted from the broth (undissociated *n*-caproic acid was ∼0.3 mM), the *n*-caproate selectivity (*i*.*e*., product *per* substrate fed) was 42.6 ± 19.0% higher compared to serum bottles without extraction. Based on the 16S rRNA gene sequence, strain 7D4C2 is most closely related to the isolates *Caproicibacter fermentans* (99.5%) and *Caproiciproducens galactitolivorans* (94.7%), which are chain-elongating bacteria that are also capable of lactate production. Whole-genome analyses indicate that strain 7D4C2, *C. fermentans*, and *C. galactitolivorans* belong to the same genus of *Caproiciproducens*. Their rBOX genes are conserved and located next to each other, forming a gene cluster, which is different than for other chain-elongating bacteria such as *Megasphaera* spp.

**Conclusions:** *Caproiciproducens* spp., comprising strain 7D4C2, *C. fermentans, C. galactitolivorans*, and several unclassified strains, are chain-elongating bacteria that encode a highly conserved rBOX gene cluster. *Caproiciproducens* sp. 7D4C2 (DSM 110548) was studied here to understand *n*-caproate production better at mildly acidic pH within microbiomes and has the additional potential as a pure-culture production strain to convert sugars into *n*-caproate.

## Introduction

Medium-chain carboxylates (MCCs, 6-12 carbon atoms) are precursors to liquid fuels (Levy et al., 1981). Production of MCCs is, therefore, of great interest to biotechnology as a production platform for large volumes, especially since the substrate can be organic wastes or wastewater as part of the circular economy. MCCs are much easier to separate from the culture broth compared to short-chain carboxylates (SCCs, 2-5 carbon atoms) due to their hydrophobic carbon chains (Angenent et al., 2016; Levy et al., 1981; Xu et al., 2015). Besides their use for fuel production, MCCs are also feedstocks in the chemical, pharmaceutical, food, and agricultural industries for the manufacture of a wide variety of products (Desbois, 2012; Harvey & Meylemans, 2014; Kenealy et al., 1995; Levy et al., 1981). Moreover, MCCs are used for food preservation and sanitation due to their antimicrobial properties at low pH (Harroff et al., 2017).

Carboxylates exist in an undissociated (carboxylic acid) and dissociated form (conjugate base, or carboxylate, plus a proton), depending on the pH of the bioreactor broth. At mildly acidic pH, specifically below the pKa (∼4.9), the carboxylic acid is in the undissociated form. At pH values higher than the pKa, the acid dissociates and releases one proton, forming the conjugate base. The undissociated form of a carboxylate (*i*.*e*., the carboxylic acid) is hydrophobic, which is essential for separation, but it is also lipophilic and crosses the microbial cell wall, creating antimicrobial properties. Inside the cell, where the pH is higher than in the bioreactor broth, the acid dissociates. As the conjugate base is lipophobic, it accumulates inside the cell, resulting in microbial inhibition (Russell, 1992). Based on this, *n*-caproate, which is a 6-carbon MCC (here referred to as the total of dissociated and undissociated forms), is toxic to microbes at pH values near its pKa (Agler et al., 2012a; Ge et al., 2015).

Chain-elongating bacteria produce MCCs *via* the reverse β-oxidation (rBOX) pathway. In this strictly anaerobic process, electron donors, such as fructose, sucrose, lactate, or ethanol, are oxidized into several acetyl-CoA molecules (2 carbons each). A certain fraction of these molecules is converted to produce acetate and energy. The other fraction of the acetyl-CoA molecules is used to elongate acetate or other SCCs (electron acceptors) in a cyclic process where two carbons are added at a time (**Figure 1**). In this manner, acetate (2 carbons) is first elongated to *n*-butyrate (4 carbons) and then to *n*-caproate (6 carbons). In some cases, *n*-caprylate (8 carbons) is produced (Kucek et al., 2016a; Kucek et al., 2016b; Spirito et al., 2014). When propionate is the electron acceptor, *n*-valerate (5 carbons) and *n*-heptanoate (7 carbons) are produced (Jeon et al., 2016). However, electron donors can also be used solely to produce MCCs (Jeon et al., 2010). The key enzymes involved in the rBOX pathway are thiolase (Thl; also named acetyl-CoA C-acetyltransferase), 3-hydroxybutyryl-CoA dehydrogenase (HBD), crotonase (Crt; also named 3-hydroxybutyryl-CoA dehydratase), acyl-CoA dehydrogenase (ACDH), electron transport flavoprotein (ETF), and acetate-CoA transferase (ACT) (**Figure 1**).

**Figure 1.**
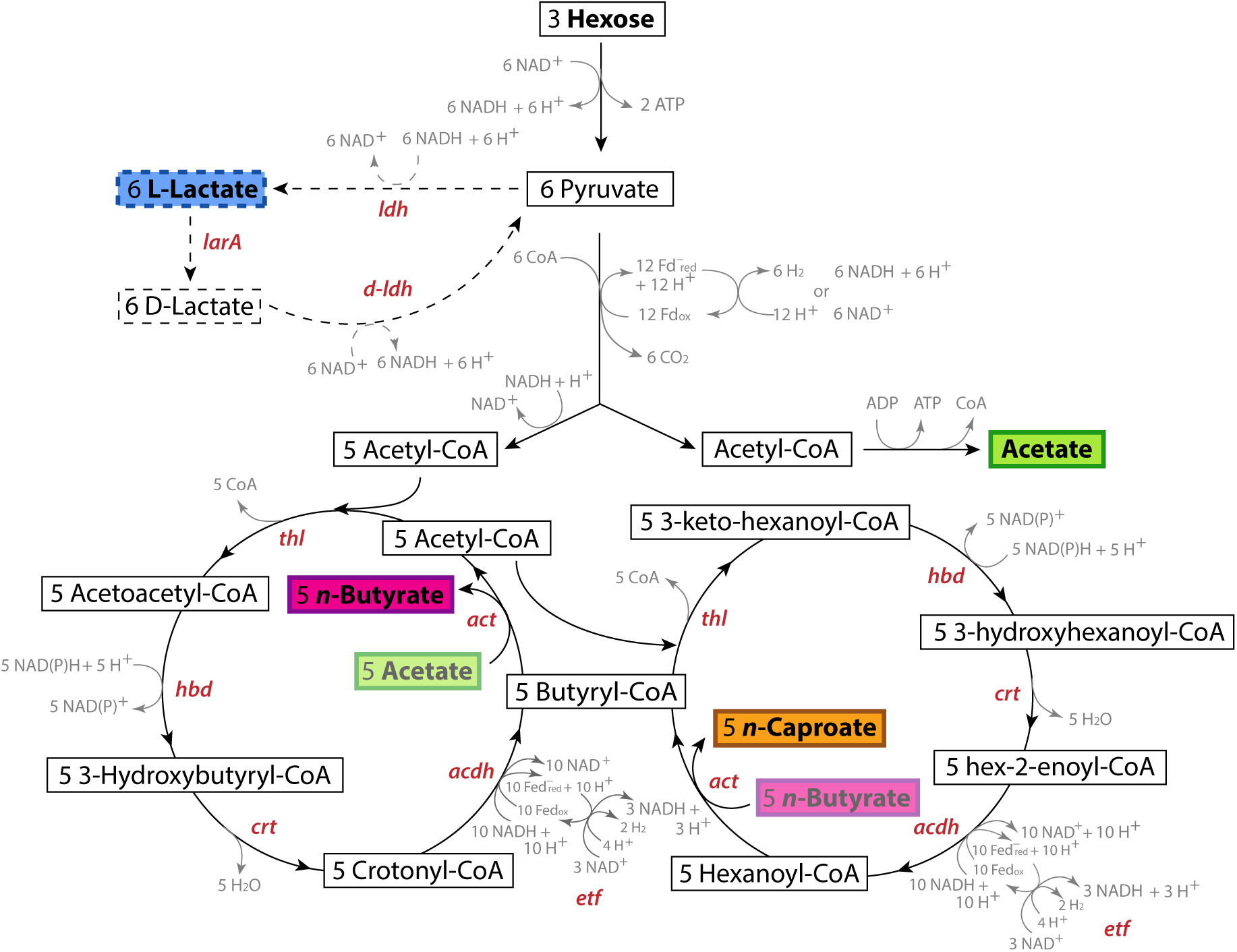
Pathways and genes involved in the conversion of hexoses into lactate and the conversion of these substrates into *n*-caproate *via* the reverse β-oxidation (rBOX) pathway. The first cycle of the rBOX pathway involves the conversion of the acetate produced by one acetyl-CoA molecule into *n*-butyrate. The second cycle involves the conversion of this *n*-butyrate into *n*-caproate *via* the butyryl-CoA produced in the first cycle and an acetyl-CoA molecule. The genes that code for the enzymes catalyzing the production of lactate and its conversion into pyruvate and each reaction of the rBOX pathway are shown for each reaction. rBOX genes: *thl*, thiolase (acetyl-CoA C-acetyltransferase); *hbd*, 3-hydroxybutyryl-CoA dehydrogenase; *crt*, crotonase (3-hydroxybutyryl-CoA dehydratase); *acdh*, Acyl-CoA dehydrogenase; *etf*, Electron transport flavoprotein; *act*, acetate-CoA transferase. Lactate production gene: *ldh, L-ldh*, L-lactate dehydrogenase. Lactate consumption genes: *larA*, lactate racemase; *D-ldh*: D-lactate dehydrogenase.

Open cultures (*e*.*g*., reactor microbiomes) have been used to generate MCCs at high rates from various synthetic feeds and industrial and agricultural wastewaters, which are rich in carbon and electron equivalents such as sugar-rich and lactate-rich effluents (Contreras-Dávila et al., 2020; Duber et al., 2018; Kucek et al., 2016a; Xu et al., 2018). These bioreactors are operated: (1) at neutral pH to circumvent the accumulation of the undissociated form of the carboxylates, or (2) at mildly acidic pH (5.0-5.5) with in-line MCC extraction to recover the carboxylate product and to prevent product inhibition. The operation of bioreactors at mildly acidic pH values has the advantage of facilitating the extraction of MCCs from the culture broth because, at these pH values, MCCs have a low maximum solubility (Xu et al., 2015). Also, the low pH in open-culture bioreactors inhibits acetoclastic methanogenesis, which would be the main, but unwanted, electron shunting mechanism in reactors operated at neutral pH (Ge et al., 2015).

To increase the likelihood that MCC production in bioreactors with in-line extraction becomes an economic proposition as a biotechnology production platform, it is essential to study chain-elongating bacteria that thrive under mildly acidic conditions. A few chain-elongating bacteria have been isolated. *Clostridium kluyveri* is the most studied chain-elongating bacterium and known to utilize ethanol as the primary electron donor (Angenent et al., 2016). Other well-studied chain-elongating bacteria use carbohydrates (*e*.*g*., *Caproiciproducens galactitolivorans, Megasphaera hexanoica, Megasphaera elsdenii, Megasphaera indica*,) or lactate (*e*.*g*., *Ruminococcaceae* bacterium CPB6 and *M. elsdenii*) as electron donors (Felicity A. Roddick & Britx, 1997; Jeon et al., 2016; Lanjekar et al., 2014; Marounek et al., 1989; Zhu et al., 2017).

Recently, *Caproicibacter fermentans*, which is an *n*-caproate producer from carbohydrates, was isolated (Flaiz et al., 2020). While open cultures can effectively perform chain elongation at mildly acidic pH conditions with in-line MCCs extraction, strain CPB6 and *C. fermentans* are the only known chain-elongating bacteria that can satisfactorily produce MCCs at mildly acidic pH levels (Flaiz et al., 2020; Zhu et al., 2017).

Whole-genome analyses combined with laboratory experiments are a powerful approach to study chain-elongating bacteria. While whole-genome alignments are necessary to assign taxonomy to novel microbes, the presence and location of genes give insights into their metabolism. The main objective of this work was to isolate and study the metabolism of a chain-elongating bacterium that thrives at mildly acidic pH (>4.5). To consider its potential application in bioreactors that are aimed at MCC production, we identified the environmental conditions that enhanced its *n*-caproate production. We sequenced and assembled its whole genome and compared it to other bacteria to assign taxonomy. We focused our comparisons on its closest isolated relatives *C. fermentans* (99.5% similar based on the 16S rRNA gene sequence) and *C. galactitolivorans* (94.71%), and also on unclassified strains. Moreover, we studied the genes encoding rBOX proteins (rBOX genes) and compared them to those in: (1) close relatives; (2) bacteria with similar rBOX genes; and (3) known-chain-elongating bacteria.

## Results

### Strain 7D4C2 is a chain-elongating bacterium that converts sugars into *n*-caproate, lactate, and H_2_ at mildly acidic pH

We cryogenically preserved a sample from an open-culture, chain-elongating bioreactor that was operated at a pH of 5.5 and 30°C and fed with ethanol and acetate in our previous laboratory at Cornell University in Ithaca, NY, USA (Spirito, Angenent, *et al*., unpublished work). We revived the sample with ethanol (40 mM), acetate (4 mM), *n*-caproate (4 mM), and *n*-caprylate (4 mM) in basal medium that was buffered with 91.5 mM MES and supplemented with 0.05% w/v yeast extract and vitamins (**Additional File 1: Figure S1**). To isolate chain-elongating bacteria, we serially diluted the culture and picked single colonies (pH 5.2, 30°C).

Next, the selected colonies were cultured in a liquid medium and further diluted for purification. Since this liquid culture did not consume ethanol, we continued the purification process with fructose as the primary electron donor. The high concentration of MES, the mildly acidic pH (5.2), as well as the added fructose and electron acceptors (*n*-butyrate, *n*-caproate, and *n*-caprylate), inflicted strong selective pressures that allowed the relatively fast isolation (**Additional File 1: Figure S1**). Ultimately, the isolate that produced *n*-caproate and showed 100% purity is referred to as strain 7D4C2 (DSM 110548). Strain 7D4C2 is a Gram-positive bacterium (**Additional File 1: Figure S2**) and rod-shaped (**Figure 2A**,**B**), which produces lactate, acetate, *n*-butyrate, *n*-caproate, biomass, and H_2_ from hexoses at a pH of 5.5 (**Figure 2C-E**). CO_2_ is also produced (data not shown).

**Figure 2.**
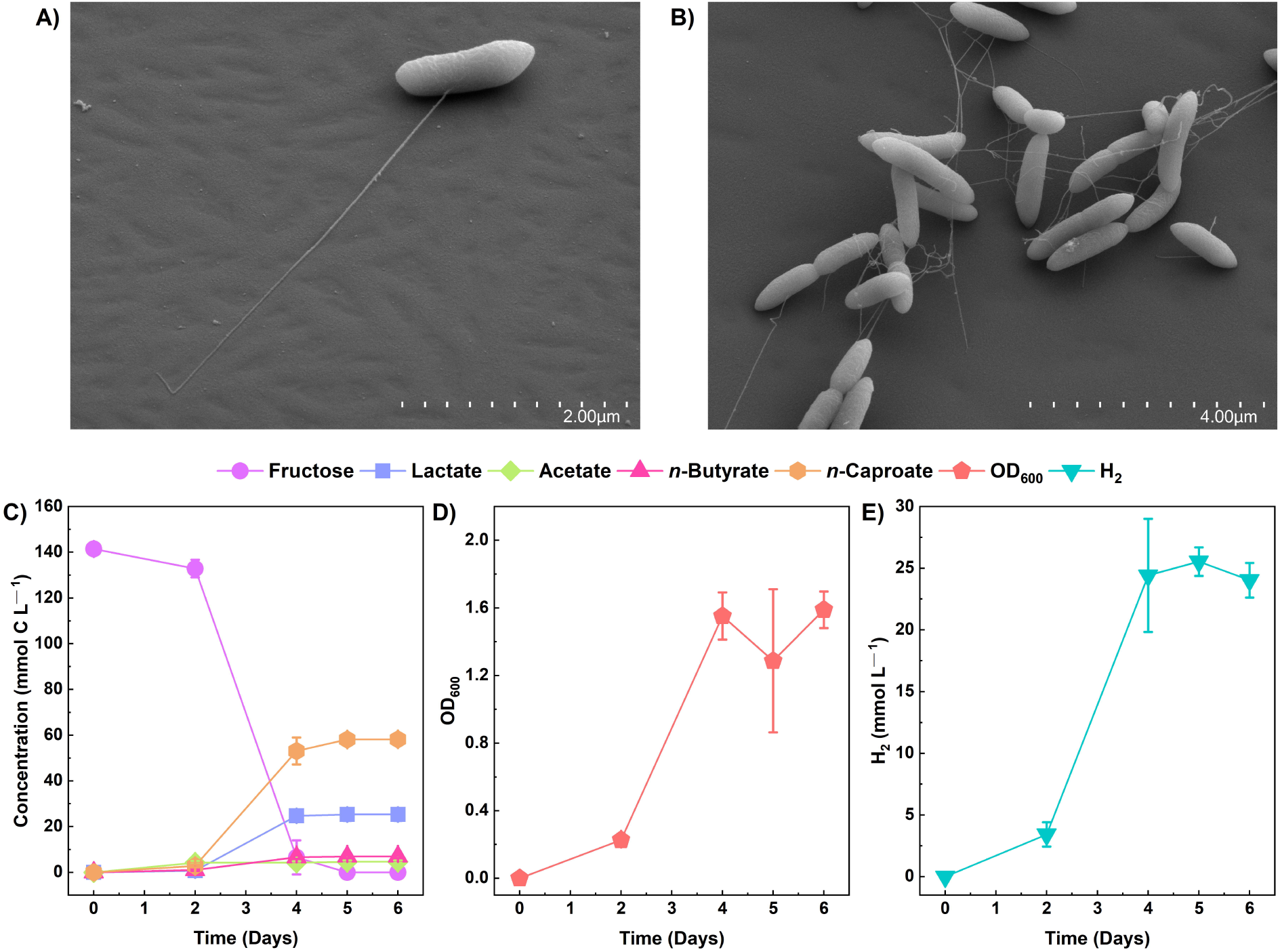
Growth of strain 7D4C2 with fructose at pH 5.5 and 30°C: A-B) scanning electron micrographs of strain 7D4C2; C) fructose conversion into *n*-caproate and lactate; D) growth measured by OD_600_; and C) H_2_ production. Error bars represent one standard deviation among triplicate cultures.

### The presence of different electron acceptors from 2 to 6 carbons influenced chain elongation by strain 7D4C2

Short-chain carboxylates are commonly used as electron acceptors in chain elongation (Jeon et al., 2016; Wang et al., 2018). To study whether strain 7D4C2 was capable of utilizing even- and odd-chain electron acceptors, we grew the isolate at a temperature of 30°C and a pH of 5.5 with fructose (146.4 ± 10.3 mmol C L^−1^) and different carboxylates (108.2 ± 8.0 mmol C L^−1^) from 2 to 6 carbons (*i*.*e*., acetate, propionate, *n*-butyrate, *n*-valerate, and *n*-caproate) in separate serum bottles. For the control (fructose without an electron acceptor), strain 7D4C2 achieved a final average concentration of 6.9 ± 0.6 mmol C L^−1^ for *n*-butyrate and 57.5 ± 2.4 mmol C L^−1^ for *n*-caproate (**Figure 3A,B**), with an *n*-caproate specificity of 60.9 ± 1.5% (*i*.*e*., products *per* other carboxylates produced) (**Additional File 1: Table S1)**. The presence of electron acceptors influenced the metabolism of strain 7D4C2. For acetate as the electron acceptor (13.8 ± 8.1% uptake), the final average *n*-butyrate concentration was higher than the control (38.7 ± 7.2 mmol C L^−1^), while the *n*-caproate concentration was lower (40.3 ± 15.4 mmol C L^−1^), with an *n*-caproate specificity of 44.1 ± 5.9% (**Figure 3A,C, Additional File 1: Table S1**). For propionate as the electron acceptor, the 47.1 ± 1.7% uptake changed the metabolism from *n*-caproate to *n*-valerate production (compared to the control serum bottles) to reach a final average *n*-valerate concentration of 76.5 ± 0.4 mmol C L^−1^, although with a longer lag phase for fructose uptake (**Figure 3A,D**). This resulted in an *n*-caproate specificity of only 2.79 ± 0.5% (**Additional File 1: Table S1**). Strain 7D4C2 achieved a higher *n*-caproate concentration for *n*-butyrate as the electron acceptor (53.3 ± 1.1% uptake) than for the control and the rest of carboxylates as electron acceptors, resulting in a total average concentration of 125.5 ± 1.9 mmol C L^−1^ and an *n*-caproate specificity of 83.1 ± 44% (**Figure 3A, Additional File 1: Table S1**). Previous studies with other chain-elongating bacteria have also observed the highest *n*-caproate specificities with *n*-butyrate (Jeon et al., 2016; Zhu et al., 2017). Moreover, the mmol-C ratio of produced *n*-caproate to lactate was higher at 20:1 for the serum bottles with *n*-butyrate than at 2:1 for the control (**Figure 3A,E, Additional File 1: Table S1**). For *n*-valerate as the electron acceptor (10.1 ± 0.7% uptake), the final average lactate concentration was higher than the rest of the serum bottles (46.2 ± 3.2 mmol C L^−1^), and equivalent to the final average *n*-caproate concentration (44.3 ± 5.3 mmol C L^−1^), with an *n*-caproate specificity of 41.4 ± 3.3% (**Figure 3A,F, Additional File 1: Table S1**). We do not completely understand the reasons for these shifts in metabolism but know from theoretical calculations that the ratio of electron donor and electron acceptor has a large thermodynamic effect on product formation (Angenent et al., 2016). Lastly, for *n*-caproate as the electron acceptor, the initial total concentration of 102.4 ± 0.5 mmol C L^−1^ resulted in an undissociated *n*-caproic acid concentration of ∼19.8 mmol C L^−1^ (∼3.3 mM) at a pH value of 5.5, which completely inhibited the metabolism of strain 7D4C2 (**Figure 3A,G, Additional File 1: Table S1)**.

**Figure 3.**
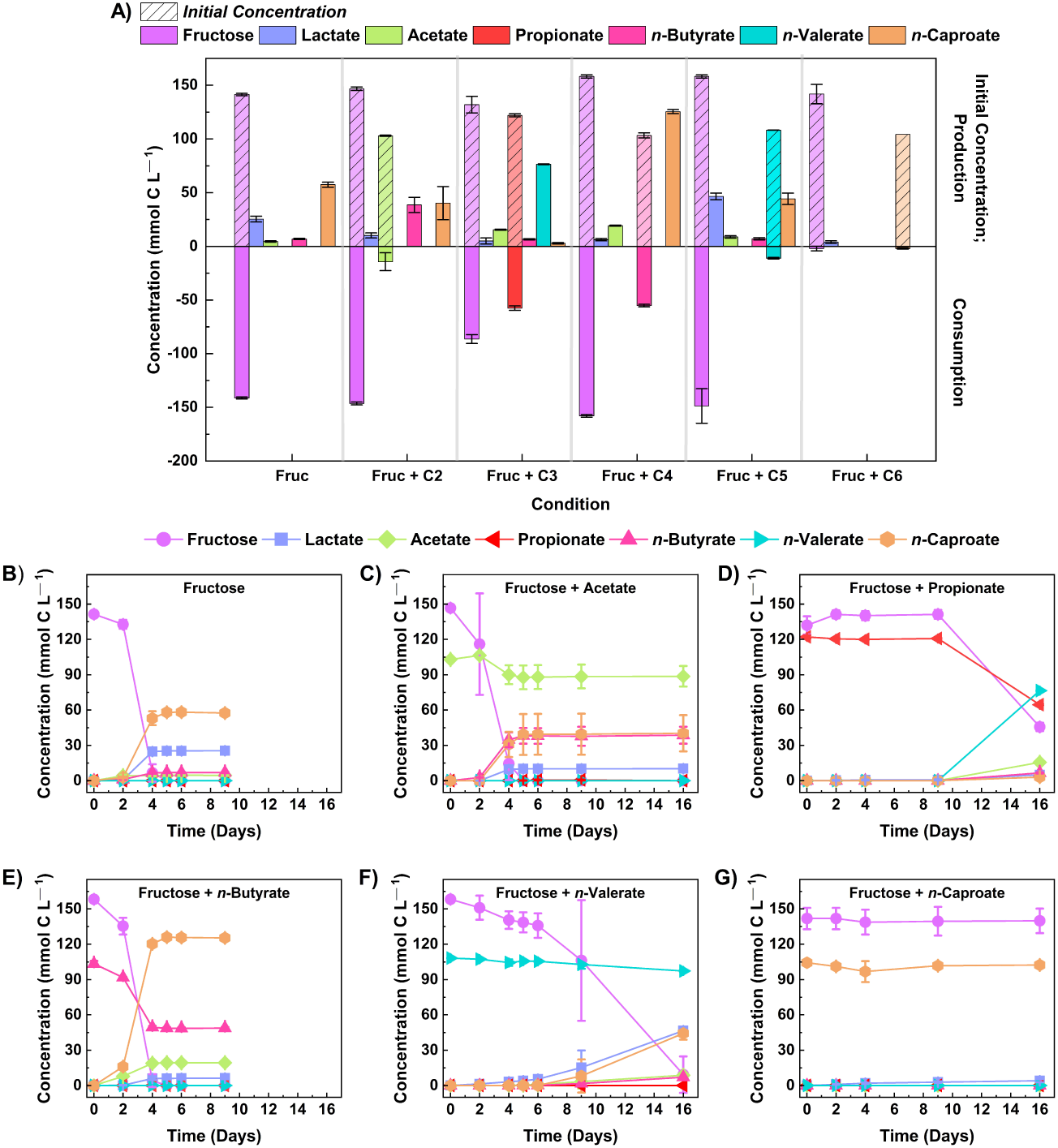
Comparison of lactate and MCCs (*i*.*e*., *n*-valerate and *n*-caproate) produced by strain 7D4C2 from fructose and different electron acceptors (C2→ C6): **A)** comparison of final products and fructose and electron donor consumption among experiments; and **B-G)** fructose, electron acceptor, and products concentrations throughout the culturing period for each electron acceptor (acetate, propionate, *n*-butyrate, *n*-valerate, and *n*-caproate, respectively). Fruc: fructose; C2: acetate; C3: propionate; C4: *n*-butyrate; C5: *n*-valerate; and C6: *n*-caproate. The initial fructose concentration was 146.4 ± 10.3 mmol C L^−1^ and the concentration of the electron acceptors was 108.2 ± 8.0 mmol C L^−1^. The pH value of the test was 5.5 ± 0.02. Error bars represent one standard deviation among triplicate cultures.

### The specificity of *n*-caproate production was higher at mildly acidic pH values while that of lactate was higher at alkaline pH levels

Next, we investigated lactate and *n*-caproate production of strain 7D4C2 at a pH gradient: from mildly acidic to alkaline pH levels. For this, we cultured strain 7D4C2 at 30°C with a mixture of fructose (148.2 ± 3.2 mmol C L^−1^) and *n*-butyrate (112.2 ± 6.3 mmol C L^−1^) as the substrate at different initial pH values from 4.5 to 9.0 (**Figure 4**). We did not manually adjust the pH during the culture period, but we strongly buffered the serum bottles with 91.5 mM MES. The initial mildly acidic pH values from 4.5 to 5.5 favored the mmol-C ratio of produced *n*-caproate to lactate (lactate below detection at a pH value of 4.5 and 13:1 mmol C L^−1^ at a pH value of 5.5), with final average *n*-caproate concentrations of 93.2 to 146.7 mmol C L^−1^ (**Figure 4A**). The average *n*-caproate specificities for pH 4.5 to 5.2 were ∼90%, but the specificity decreased to ∼83% for the pH 5.5 condition (**Additional File 1: Table S2**). At initial pH values higher than 6.0, the mmol-C ratio of produced *n*-caproate to lactate gradually decreased to 0.4:1 at a pH value of 9.0. Strain 7D4C2 achieved a maximum average lactate concentration of 103.0 mmol C L^−1^ at a pH of 9.0 (**Additional File 1: Table S2**). In addition, strain 7D4C2 metabolized less and less *n*-butyrate across the increasing pH gradient (**Figure 4A**). Together, the changes in metabolism across the alkaline pH values led to a decrease in the final average *n*-caproate concentration from <76.0 to ∼36.0 mmol C L^−1^ for pH 7.0 to 9.0 (**Figure 4A**), resulting in a decrease in specificity from 37 to 23% (**Additional File 1: Table S2**). The H_2_ production in mmol L^-1^ did not follow the exact same trend of *n*-caproate specificity, but it was the highest at the low pH values of 5.2 and 5.5 (**Figure 4B**). We also cultured strain 7D4C2 at an initial pH of 10.0, but it did not grow (data not shown).

**Figure 4.**
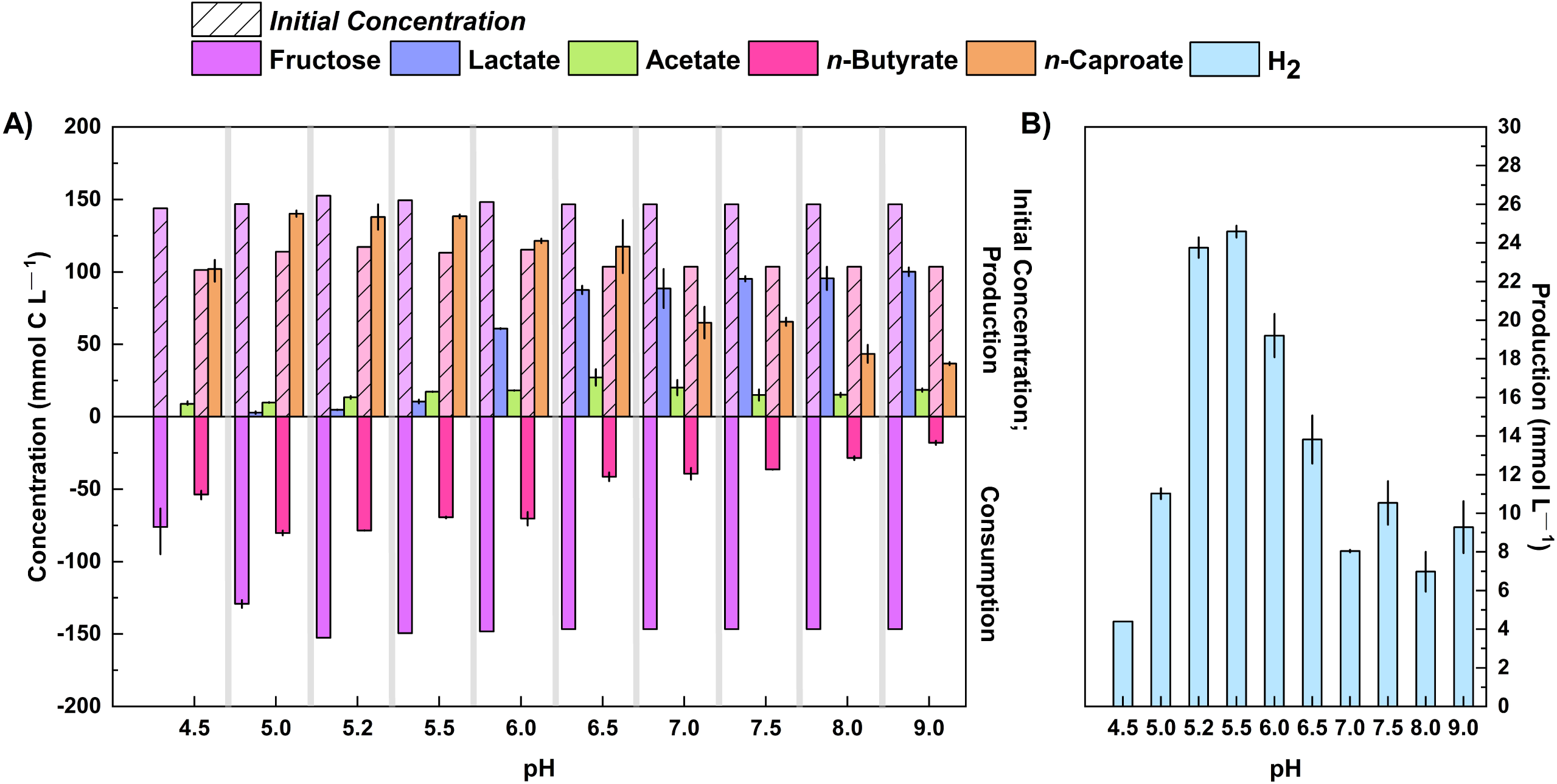
Production of lactate, *n*-caproate, and H_2_ by strain 7D4C2 across a wide pH range (4.5 to 9.0*): **A)** comparison of final products (lactate, acetate, and *n*-caproate) and fructose and *n*-butyrate consumption among experiments at different pH values; and **B)** comparison of final H_2_ production among experiments at different pH values. Bars represent minimum and maximum values between duplicate cultures. The initial concentrations of fructose and *n*-butyrate are shown in transparency with a lined pattern. *Initial pH values in an MES-buffered system. The lag phase at pH values of 4.5 and 5.0 was slower than the rest of the experiments (see **Figure S3**).

### The optimum pH and temperature for *n*-caproate production differed for the growth rate

As discussed in the previous section, strain 7D4C2 achieved the highest *n*-caproate specificity at mildly acidic pH values (4.5 – 5.2). However, at a pH of 4.5 and 5.0, the bacterium grew with an extended lag phase compared to the pH values 5.2 and 5.5 (**Additional File 1: Figures S3A**,**B and S4A**). Based on the high *n*-caproate specificity (∼88.3%) and concentration (∼3.1 mmol C L^−1^) in combination with a high growth rate (0.5 d^−1^), the optimum pH value for improved *n*-caproate production was 5.2 (**Additional File 1: Table S2**). However, this pH value differed from the optimum pH value for growth, which was 6.0. At an initial pH of 6.0, the H_2_ production rate, growth rate (1.3 d^−1^) (**Additional File 1: Figure S3A-D**), and fructose consumption rate (37.0 mmol C L^−1^ d^−1^; **Additional File 1: Figure S4A-I, Table S2**) were the highest for this study, but the strain produced an equivalent mixture of *n*-caproate and lactate (2:1 mmol C L^−1^ in **Additional File 1: Table S2**), resulting in a lower *n*-caproate specificity than at a pH of 5.2.

Similar to the experiment with different pH values, we investigated the optimum temperature for *n*-caproate production and growth with strain 7D4C2. For this, we grew the isolate with fructose and *n*-butyrate at different temperatures, ranging from 22.5°C to 50°C, and at a pH 6.0 (the optimum pH for growth) in separate serum bottles. We found that strain 7D4C2 achieved a maximum *n*-caproate specificity of ∼67% at a temperature of 30°C (∼107 mmol C L^−1^ in **Additional File 1: Table S2**). However, similar to the pH optimum, the optimum temperature for *n*-caproate production differed for the growth rate, which was 37°C and 42°C. At these temperatures, the fructose consumption rate was 45.5 mmol C L^−1^ d^−1^, compared to 27.3 mmol C L^−1^ d^−1^ at 30°C, and the H_2_ production rate was the highest (**Additional File 1: Figure S3E-F, Figure S4J-N**).

### Product extraction increased the *n*-caproate selectivity at a pH of 5.2

Bioreactors that were operated at mildly acidic pH with in-line product extraction have shown promising MCC production rates and yields (Agler et al., 2014; Ge et al., 2015; Kucek et al., 2016a; Kucek et al., 2016b; Spirito et al., 2018). Accordingly, we tested whether strain 7D4C2 could achieve a higher *n*-caproate selectivity (*i*.*e*., product *per* substrate fed) when the MCC was extracted during growth, avoiding the toxicity of the undissociated form at mildly acidic pH. For this, we cultured the bacterium with fructose (314.1 ± 2.1 mmol C L^−1^) and *n*-butyrate (101.3 ± 3.2 mmol C L^−1^) as substrates, with product extraction and without product extraction (control) at a pH level of 5.2 and a temperature of 30°C. With the extraction of *n*-caproate, the average concentration of the undissociated MCC in the culture medium remained low at 0.3 ± 0.16 mM, while *n*-caproate production continued until all fructose was depleted by day 7 (**Figure 5D, Additional File 1: Figure S5B**). Without extraction, strain 7D4C2 reached the stationary growth phase by day 5 with substrate left over due to inhibition at an undissociated *n*-caproic acid concentration of 4.8 mM (**Figure 5A-C, Additional File 1: Figure S5A**). As a result, product extraction of *n*-caproate resulted in a 42.6 ± 19.0% higher *n*-caproate selectivity than the control without extraction (*i*.*e*., 62.9 ± 39.7 mmol C L^−1^ more *n*-caproate produced). These results indicate that *Caproiciproducen*s sp. 7D4C2 has the potential as a chain-elongating production bacterium when extraction is desired for sugars as the electron donor.

**Figure 5.**
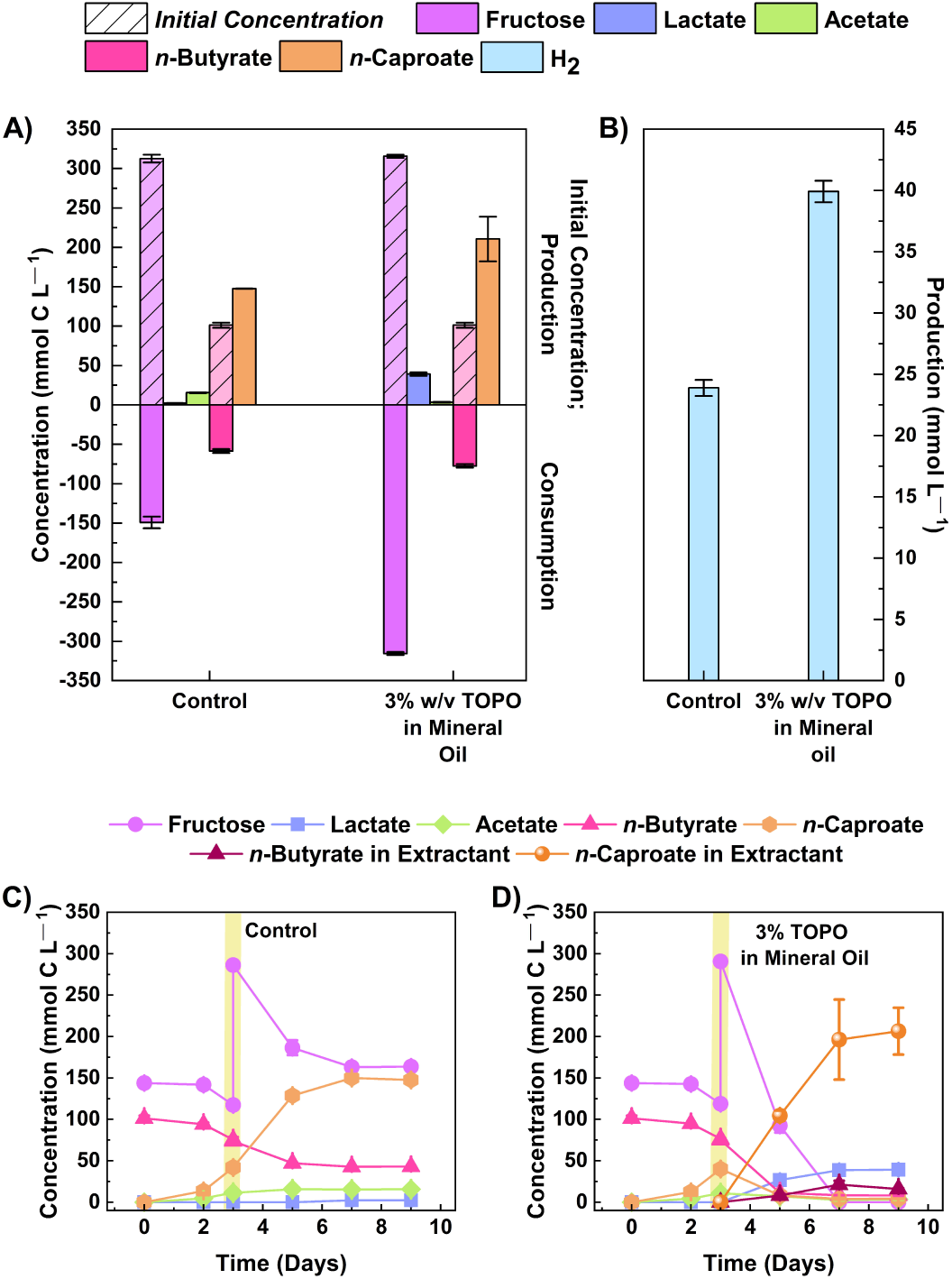
Comparison of *n*-caproate production by strain 7D4C2 with and without product extraction: **A)** comparison of final products (lactate, acetate, and *n*-caproate) and fructose and *n*-butyrate consumption between experiments with and without mineral oil and 3% w/v TOPO to extract products; **B**) comparison of of final H_2_ production between experiments with and without product extraction; **C-D)** fructose, *n*-butyrate, and products concentrations throughout the culturing period for the experiments without (C) and with product extraction (D). Vertical yellow lines represent the time-point were the fructose was increased (to increase *n*-caproate production) and an equal volume of mineral oil with 3% w/v TOPO was added. Error bars represent one standard deviation among triplicate culture.

### Strain 7D4C2 is closely related to unclassified *Clostridium* sp. W14A, *C. fermentans*, unclassified *Caproiciproducens* sp. NJN-50, and *C. galactitolivorans*

To assign taxonomy to strain 7D4C2, we sequenced its genome *via* long-read Nanopore sequencing. We obtained 117,171 reads, with an average length of 4,211 bp (N50 of 8772 bp) and a total size of 486Mb. The genome assembly resulted in a single, circular, and closed chromosome with a full length of 3,947,358 bp and a GC content of 51.6%. It was annotated with 3633 protein-coding genes (CDS), 13 rRNA genes (five 5S rRNA genes, four 16S rRNA genes, and four 23S rRNA genes), 60 tRNA genes, 4 ncRNA genes, 1 tmRNA gene, and 203 pseudogenes (154 frameshifted genes). The assembly was 97.85% complete and 1.68% contaminated, according to CheckM (Parks et al., 2015). We aligned the whole genome against the NCBI-nt database. The strain is most similar to four known bacteria: (1) unclassified *Clostridium* sp. W14A (average nucleotide identity, ANI = 97.64; 82.28% aligned bases); (2) *C. fermentans* EA1 (ANI = 97.34; 81.20% aligned bases); (3) unclassified *Caproiciproducens* sp. NJN-50 (ANI = 78.52; 44.36% aligned bases); and (4) *C. galactitolivorans* BS-1 (ANI = 69.65; 28.22% aligned bases) (**Additional File 1: Table S3**). The ANI values for the genome comparison of strain 7D4C2 with *Clostridium* sp. W14A and *C. fermentans* were higher than the cut-off value of 95 – 96% to define a novel species (∼97.5%; **Additional File 1: Table S3**) (Richter & Rosselló-Móra, 2009; Yarza et al., 2014), which indicates that these three bacteria represent different strains of the same species.

To investigate further, we also compared the 16S rRNA gene sequences from strain 7D4C2 with closely related bacteria. We identified four different 16S rRNA gene sequences (1517 – 1524 bp) in the genome of strain 7D4C2, which were 99.03% similar among them. To calculate phylogenetic distances with the other four bacteria, we aligned their 16S rRNA gene sequences (Project ID PRJNA615378) and the Sanger assembly for one of the 16S rRNA gene sequences in strain 7D4C2 (1287 bp, NCBI <u>MT056029</u>) against the NCBI-nt (National Center for Biotechnology Information; ftp://ftp.ncbi.nlm.nih.gov/blast/db/FASTA/ - accessed January 2020). Since the 16S rRNA gene sequence for *Clostridium* sp. W14A was not publicly available, we annotated the genome for W14A and extracted the 16S rRNA gene sequence. The high-to-low similarities of the 16S rRNA gene sequence for strain 7D4C2 to the four bacteria were in the same order as when the genome alignment was compared: (1) unclassified *Clostridium* sp.

W14A (100% similarity to the entire 16S rRNA gene sequence); (2) *C. fermentans* (99.51 ± 0.25% similarity); (3) unclassified *Caproiciproducens* sp. NJN-50 (97.72 ± 0.31%); and (4) *C. galactitolivorans* (94.71 ± 0.35% similarity) (**Additional File 1: Figure S6**). A cross comparison for *Clostridium* sp. W14A and *C. fermentans* to *C. galactitolivorans* showed us a 94.83% similarity between *Clostridium* sp. W14A and *C. galactitolivorans*, and a 94.90% similarity between *C. fermentans* and *C. galactitolivorans*, which is slightly outside the quantitative window to group all four strains within a single genus (Yarza et al., 2014). Thus, based on both the genome alignment and 16S rRNA gene sequence comparisons, strain 7D4C2 and its four closest related bacteria are not all strains of the same species, but likely they are all members of the same genus of *Caproiciproducens* spp.. This would mean that *C. fermentans* (*Caproicibacter fermentans*) would need to be re-classified as *Caproiciproducens fermentans*.

### The percentage of conserved proteins also suggest that strain 7D4C2, *C. fermentans*, and *C. galactitolivorans* belong to the same genus, but not the same species

To further study whether strain 7D4C2 and its closest related bacteria are members of a single species or a single genus, we calculated the percentage of conserved proteins (POCP) for strain 7D4C2, *C. fermentans, C. galactitolivorans*, and their closely related unclassified strains (*i*.*e*., *Clostridium* sp. W14A, *Caproiciproducens* sp. NJN-50, and *Clostridium* sp. KNHs216). Besides *Clostridium* sp. KNHs216, we also included additional selected species from the Clostridiales (according to the NCBI Taxonomy Database; heterotypic synonym of Eubacteriales) for this analysis (those with the highest ANI values with strain 7D4C2, **Additional File 1: Table S3**). Qin et al. 2014 have suggested that species within the same genus share at least half of their proteins, and therefore their pairwise POCP values are higher than 50% within a clade (Qin et al., 2014). As anticipated from the above results, the pairwise POCP values among strain 7D4C2, *Clostridium* sp. W14A, and *C. fermentans* were high (83.4 – 87.5%). These three bacteria formed a clade with pairwise POCP values higher than 51.7% with *C. galactitolivorans* and the closely related unclassified strains (*i*.*e*., *Caproiciproducens* sp. NJN-50 and *Clostridium* sp. KNHs216), suggesting that all these bacteria belong to the same genus (Qin et al., 2014) (**Figure 6A**). However, strain 7D4C2, *Clostridium* sp. W14A, *C. fermentans*, and *Caproiciproducens* sp. NJN-50 (POCP: 61.3 - 87.5%) separated into a different sub-clade from *C. galactitolivorans* and *Clostridium* sp. KNHs216 (POCP: 59.7%) (**Figure 6A**). The former sub-clade with strain 7D4C2 separated again into two clades with *Caproiciproducens* sp. NJN-50 as the sole strain. Strain 7D4C2, *Clostridium* sp. W14A, and *C. fermentans* are very similar strains and form a separate species based on this analysis and the genome alignment comparison.

**Figure 6.**
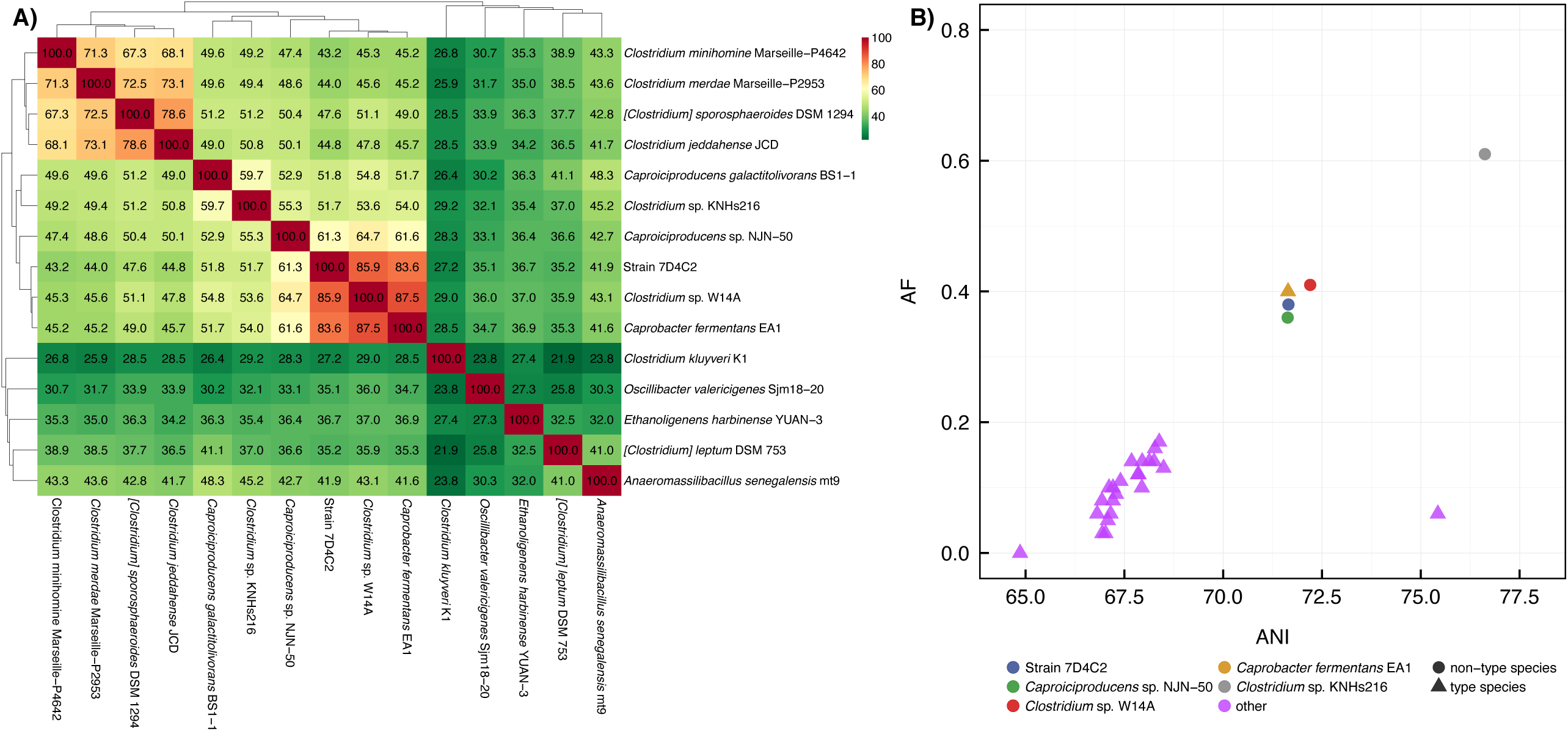
Whole-genome relatedness analyses: **A)** percentage of conserved proteins (POCP) pairwise values between selected species within the Clostridiales (heterotypic synonym of Eubacteriales). The higher the POCP value (green to red), the closer their evolutionary and genetic distance (Qin et al. 2014). The POCP analysis was performed with genomes publicly available at the NCBI; and **B)** pairwise ANI (average nucleotide identity) and AF (alignment fraction) values between *C. galactitolivorans* BS-1 and type species (*i*.*e*., first accepted species of a genus) of the *Ruminococcaceae* family (heterotypic synonym of *Oscillibacteriaceae*) (magenta), *C. fermentans* EA1 (gold), strain 7D4C2 (red), and three closely related unclassified species (in blue, green, and cyan). The validly published type species information was retrieved from The NamesforLife Database, as suggested in Barco *et al*., 2020.

In addition, we followed the approach that was suggested by Barco et al. (2020) to demarcate genera based on the relation between genome indices and the distinction of type- and non-type species. We used the average nucleotide identity of protein-coding genes and the genome aligned fraction (AF) as considered indices (Barco et al., 2020). For this analysis, we chose *C. galactitolivorans* as a reference bacterium and compared its genome relatedness index (the relation between ANI and AF) to strain 7D4C2, *C. fermentans*, and its three closely related unclassified strains (*i*.*e*., *Clostridium* sp. W14A, *Caproiciproducens* sp. NJN-50, and *Clostridium* sp. KNHs216), as well as the type species of each genus within the family *Ruminococcaceae* (according to the NCBI Taxonomy Database; heterotypic synonym of *Oscillibacteriaceae*). Results from this analysis supported our other analyses: strain 7D4C2 clustered closely with *Clostridium* sp. W14A, *C. fermentans*, and *Caproiciproducens* sp. NJN-50 (**Figure 6B**) at higher ANI and AF values than the type species, indicating the similarity to *C. galactitolivorans*. We found still higher ANI and AF values for *Clostridium* sp. KNHs216, which indicates a closer similarity to *C. galactitolivorans* than the other four bacteria (**Figure 6B**). The clear separation between strain 7D4C2, *C. fermentans*, and the three unclassified strains from the type species within the *Ruminococcaceae* suggests that neither of these species represents a novel genus, but that they are all members of the *Caproiciproducens*. This differentiation further suggests that strain 7D4C2, *C. fermentans*, and closely related unclassified bacteria are not type-species of a novel genus within their taxonomic family, but that they are part of the *Caproiciproducens*.

### Strain 7D4C2, *C. fermentans*, and *C. galactitolivorans* belong to the same genus based on their phenotype

To further validate that strain 7D4C2, *C. fermentans*, and *C. galactitolivorans* are members of the genus *Caproiciproducens*, we cultured strain 7D4C2, *C. galactitolivorans*, and [*Clostridium*] *leptum* under similar conditions (*i*.*e*., complex medium, 37°C, pH of 7.0) and compared the products from glucose fermentation. We chose [*Clostridium*] *leptum* as our reference, because it is the closest isolate to *C. galactitolivorans* (Kim et al., 2015), and it is closely related to strain 7D4C2 (**Additional File 1: Figure S6**). We then compared our results to those reported for *C. fermentans* EA1 in Flaiz *et al*. Both strain 7D4C2 and *C. galactitolivorans* produced lactate, acetate, *n*-butyrate, *n*-caproate, and H_2_/CO_2_, although at different proportions (**Additional File 1: Figure S7**). Final average lactate and *n*-caproate concentrations in serum bottles with strain 7D4C2 were higher than in serum bottles with *C. galactitolivorans*; the lactate concentration was 10.1 ± 0.7 mmol C L^−1^ higher, and the *n*-caproate concentration was 29.1 ± 0.5 mmol C L^−1^ higher (**Additional File 1: Figure S7A**). Similarly, the final average *n*-caproate concentration in serum bottles with strain 7D4C2 was 36.1 ± 1.1 mmol C L^−1^ higher than in serum bottles with C. *galactitolivorans* in a supplemented basal medium at 37°C and a pH of 6.0 (data not shown). [*C*]. *leptum* did not produce lactate nor *n*-caproate, and only ethanol and acetate were detected in the serum bottles (**Additional File 1: Figure S7A**). All three strains produced H_2_, but H_2_ production by *C. galactitolivorans* was the highest (**Additional File 1: Figure S7B**). Similar to strain 7D4C2 and *C. galactitolivorans, C. fermentans* also produced lactate, acetate, *n*-butyrate, *n*-caproate, and H_2_/CO_2_ from hexoses (Flaiz et al., 2020).

To identify phenotypic differences between strain 7D4C2, *C. fermentants*, and *C. galactitolivorans*, we studied the carbohydrate utilization of strain 7D4C2 using the AN MicroPlate^™^ from Biolog (Hayward, CA) (**Additional File 1: Figure S8**) and we compared the results to those reported for *C. fermentans* (Flaiz et al., 2020) and *C. galactitolivorans* (Kim et al., 2015). From the seven carbohydrates compared between strain 7D4C2 and *C. fermentans*, all but glycerol (oxidized by strain 7D4C2 and *C. galactitolivorans*) and D-galactose (oxidized by *C. fermentans* and *C. galactitolivorans*) showed similar utilization (**Additional File 1: Table S4**). The carbohydrate utilization by strain 7D4C2 and *C. galactitolivorans* differed in 13 out of 30 carbohydrates compared (**Additional File 1: Table S4**). Other differential characteristics between strain 7D4C2, *C. fermentans, C. galactitolivorans*, and [*C*.] *leptum* included optimal pH and temperature and genome length (**Table 1)**.

**Table 1.** Differential characteristics of strain 7D4C2 and closely related species: **1** Strain 7D4C2; **2** *Caproicibacter fermentans* (Flaiz et al. submitted); **3** *Caproiciproducens galactitolivorans* BS-1 (Kim et al., 2015a); and **4** [*Clostridium*] *leptum* VPI T7-24-1 (Moore et al., 1976).

| Characteristic | 1* | 2 | 3 | 4 |
| --- | --- | --- | --- | --- |
| Source | Anaerobic reactor | Anaerobic reactor | Anaerobic reactor | Fecal flora |
| 16S rRNA percent identity, %<sup>a</sup> | - | 99.51 ± 0.25% | 94.71 ± 0.35 | 93.44 ± 0.14 |
| POCP, % | - | 83.4 | 51.9 | 35.0 |
| Cell shape | rod | rod | rod | rod |
| Cell size (µm) | 0.3-0.4 x 1.3-2 | 0.4-0.8 x 1-3 | 0.3-0.5 x 2-4 | 0.6-8.8 x 1.3-2.8 |
| Gram stain | + | -<sup>b</sup> | + | + |
| Spores observed | No<sup>c</sup> | No<sup>c</sup> | No<sup>c</sup> | Yes |
| Optimum pH | 6.0 | 7.0 | 6.5-7.5 | NR |
| Optimum T | 37-42 | 37 | 40 | 37 |
| Substrate / Products | glucose, fructose / H<sub>2</sub>, CO<sub>2</sub>, acetate, *n*-butyrate, *n*-caproate, lactate | fructose / H<sub>2</sub>, CO<sub>2</sub>, acetate, *n*-butyrate, *n*-caproate, lactate, ethanol | glucose*, galactitol / H<sub>2</sub>, CO<sub>2</sub>, acetate, *n*-butyrate, *n*-caproate, ethanol, lactate* | maltose / H<sub>2</sub>, acetate. Glucose* / H<sub>2</sub>, CO<sub>2</sub>, ethanol*, acetate |
| GC content, % | 51.6 | 51.25 | 48.1 | 50.2 |
| Genome length, Mbp | 3.95 | 3.9 | 2.58 | 3.27 |

\*Data from this study. NR: not reported.
<sup>a</sup>The 16S rRNA gene percent identity represents an average of the percent identities obtained from the four 16S rRNA gene sequences of strain 7D4C2 extracted from the genome (NCBI PRJNA615378) and the assembly done with Sanger Sequencing (NCBI MT056029).
<sup>b</sup>Negative staining but cell wall typical of Gram-positive bacteria.
<sup>c</sup>Spores not observed but the genome encodes one or more sporulation genes.
The genomes of *C. fermentans*, *C. galactitolivorans*, and *C. leptum* were downloaded from the NCBI (accession numbers, NZ\_VWXL000000000, SRMQ000000000 and ABCB000000000, respectively). POCP: Percentage of conserved proteins with strain 7D4C2.

In general, our work shows that strain 7D4C2 and *C. fermentans* have a similar phenotype to *C. galactitolivorans*. Therefore, also based on the ∼5% dissimilarity between their 16S rRNA gene sequences and the >51.7% shared conserved proteins, we propose that: (1) strain 7D4C2, the unclassified *Clostridium* sp. W14A, *C. fermentans*, the unclassified *Caproiciproducens* sp. NJN-50, *C. galactitolivorans*, and the unclassified *Clostridium* sp. KNHs216 belong to the genus *Caproiciproducens*; and (2) strain 7D4C2, the unclassified *Clostridium* sp. W14A, and *C. fermentans*, are very similar strains of a new species within the *Caproiciproducens*. We propose *Caproiciproducens fermentans* as the name for these three strains based on the work by Flaiz et al. (2020).

### The six rBOX genes in *Caproiciproducens* species are located next to each other, forming a gene cluster

To further study the chain-elongation metabolism of strain 7D4C2, we identified the rBOX genes (*thl, hbd, crt, acdh*, and *etf-*α and *-* β; **Figure 1**) in its genome and we compared them to those in: (1) closely related bacteria (*i*.*e*., the proposed *Caproiciproducens* species); (2) bacteria with similar rBOX genes (*i*.*e*., *Anaeromassilibacillus senegalensis, Eubacterium limosum*, and several *Clostridium* species); and (3) well known chain-elongating bacteria (*i*.*e*., *C. kluyveri, Oscillibacter valerigenes*, unclassified *Ruminococcaceae* CPB6, *M. hexanoica*, and *M. elsdenii*). The number of copies for each gene varied from 1 to 14 for the included bacteria (**Additional File 2**). The genomes of strain 7D4C2, unclassified *Clostridium* sp. W14A, and *C. fermentans* EA1 have two copies for *thl*, 2-3 copies for *acdh* and *etf-*α, three copies for *etf-*β, and one copy for *hbd* and *crt*. Differently, *Caproiciproducen*s sp. NJN-50 and *Clostridium* sp. KNHs216 encode several copies for each rBOX gene, and *C. galactitolivorans* has only one copy for each gene (**Additional File 2**). In general, the genomes of the analyzed bacteria contain multiple copies for some or all of the rBOX genes. However, *C. galactitolivorans, A. senegalensis*, and uncultured *Ruminococcaceae* CPB6 only contain a single copy (**Additional File 2**).

One copy for each of the rBOX genes (*thl, hbd, crt, acdh*, and *etf-*α and *-*β) in strain 7D4C2 are located next to each other, forming a 5,903-base pair-long gene cluster (**Figure 7A)**. We observed the same synteny of the rBOX cluster for the genomes of bacteria that are closely related to the *Caproiciproducens*. Similarly, this synteny was found for *A. senegalensis*, which is not known as a chain elongator, and *E. limosum*, which is an acetate and *n*-butyrate producer (Park et al., 2017; Roh et al., 2011), and which is capable of *n*-caproate production at high *n*-butyrate concentrations (Lindley et al., 1987) (**Figure 7A**). The arrangement of the rBOX genes varied for other bacteria. For the *Clostridium* species (*i*.*e*., *Clostridium jeddahense, Clostridium sporosphaeroides, Clostridium minihomine*, and *Clostridium merdae*), which are not known to produce *n*-caproate, the rBOX gene cluster is separated; *thl* and *hbd* form one cluster and *crt, acdh, etf-*α, and *etf-*β form a separate cluster, approximately 5 kbp away from each other and on the opposite strand (**Figure 7A, Additional File 2**). For the well-known chain-elongating bacteria *C. kluyveri* and *O. valericigenes* (an *n*-valerate producer (Lino et al., 2007)), their genomes have one copy of five rBOX genes (all but *thl*) in synteny (**Figure 7A**). The *thl* genes in these two chain-elongating bacteria are separated from the rest of the rBOX genes. The three thiolase genes in *C. kluyveri* form a separate cluster 658,054 bp away from the rBOX cluster (**Additional File 2**). In *Ruminococcaceae* bacterium CPB6, *acdh, etf-*α, and *etf-*β cluster together, while *thl, hbd*, and *crt* cluster further away (924,173 bp) from the first three genes (**Figure 7A** and **Additional File 2**). The rBOX genes of *M. hexanoica* and *M. elsdenii* are not in an apparent synteny, although those of *M. hexanoica*, except *thl*, are close to each other (**Additional File 2**). More work is required to understand whether an advantage exists for chain-elongating bacteria with a gene cluster for rBOX genes compared when these genes are located separately on the genome.

**Figure 7.**
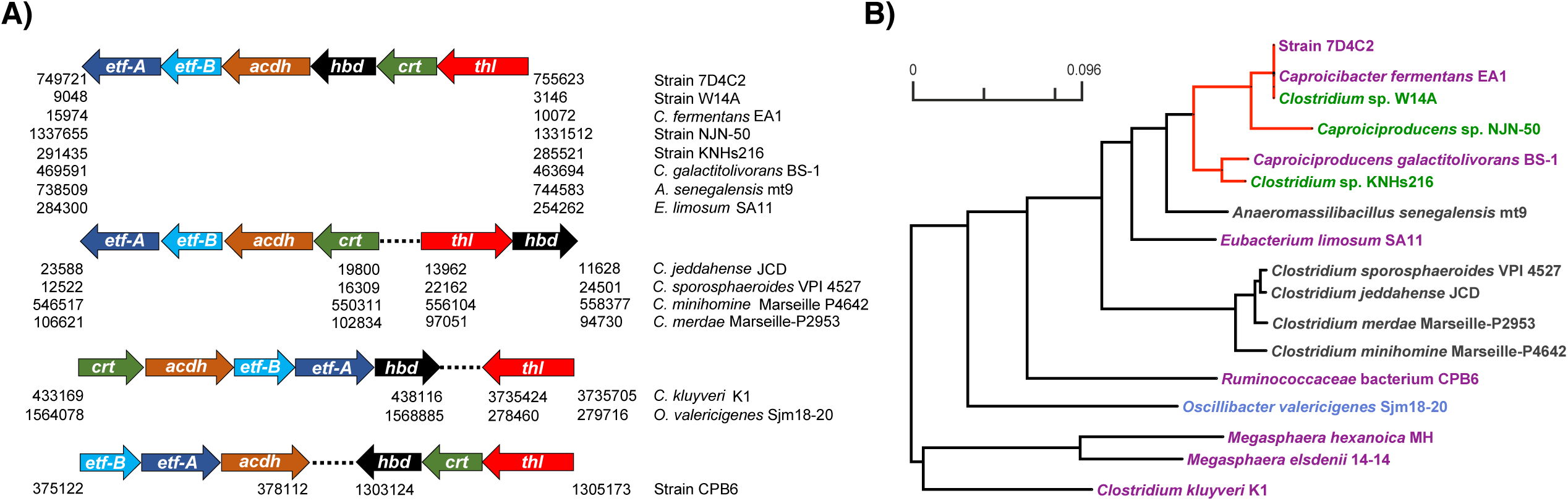
Reverse β-oxidation genes for strain 7D4C2 and bacteria with similar genes, as well as in known *n*-caproate producers: **A)** position of the rBOX genes that cluster together in these bacteria. The numbers below the arrows indicate the position (base pairs) of the genes for each bacterium on the right column; and **B)** consensus phylogenetic tree of all 6 rBOX genes that cluster together*. Red lines indicate the *Caproiciproducens* clade. Microbial names highlighted in purple denote *n*-caproate producers, in green are potential *n*-caproate producers, and in blue *n*-valerate producers. The phylogenetic distances of each of the rBOX genes in these bacteria are shown in **Figure S9**. *As the rBOX genes in the *Megasphaera* species do not cluster, for this analysis, we considered the genes most similar to strain 7D4C2.

### The rBOX genes in strain 7D4C2 are mostly similar to those in *Caproiciproducens* species and relatively distant to those in other chain-elongating bacteria

We built individual gene trees with the 6 rBOX genes and a consensus tree out of them in strain 7D4C2, closely related bacteria, bacteria with similar rBOX genes, and known chain-elongating bacteria. As the gene copies varied for different bacteria, we included in the analyses the rBOX genes that are located close to each other (forming a cluster) or that are most similar to those in strain 7D4C2 (**Additional File 2**). The analysis showed that the rBOX genes of strain 7D4C2 are identical to those of *Clostridium* sp. W14A and *C. fermentans*. In general, these genes are very similar to those of other members of the POCP clade (*i*.*e*., *Caproiciproducen*s sp. NJN-50, *C. galactitolivorans*, and *Clostridium* sp. KNHs216; **Figure 6A**) (**Figure 7B**). The rBOX genes of strain 7D4C2 are also similar to those of less closely related bacteria, such as *A. senegalensis* and *E. limosum*, but relatively distant to those of other chain-elongating bacteria (*i*.*e*., *C. kluyveri, O. valerigenes, Ruminococcaceae* bacterium CPB6, *M. hexanoica*, and *M. elsdenii*) (**Figure 7B**).

The individual gene trees showed that the phylogenetic distance between the rBOX genes of strain 7D4C2 and related bacteria varies for each gene. Nonetheless, the rBOX genes of the proposed *Caproiciproducens* spp. are often within a monophyletic clade, and are always close to each other (**Additional File 1: Figure S9**). The rBOX genes of *A. senegalensis* and *E. limosum* are phylogenetically closest to those of the *Caproiciproducens*. In the cases of *acdh* and *etf-*β, these bacteria form a cluster together with *Caproiciproducens* species. The exceptions are *thl* and *hbd* in *E. limosum*, which are distant to the *Caproiciproducens* and closer to the *Clostridium* species (**Additional File 1: Figure S9**). The lactate consumer *Ruminococcaceae* bacterium CPB6 shows an interesting pattern in the individual gene trees. In the gene trees of *thl* and *crt*, strain CPB6 clusters within the *Caproiciproducens* clade, but it is distant to these bacteria in the rest of the gene trees (**Additional File 1: Figure S9**). Because of this, in the consensus tree, strain CPB6 is relatively distant to strain 7D4C2 (**Figure7B**). In summary, the distances of the rBOX genes varied among individual gene trees, both within well-known and not known chain-elongating bacteria, showing no consensus on a particular gene being relatively more conserved in chain-elongating bacteria than other bacteria.

## Conclusions

We isolated a chain elongating bacterium (strain 7D4C2) that primarily produces *n*-caproate from carbohydrates at mildly acidic pH values (4.5-5.5). The isolate has the potential to be used in chain-elongating bioreactors that treat organic waste and are operated at mildly acidic pH with in-line product extraction. After extensive comparison of the whole-genomes of strain 7D4C2 with the isolates *C. galactitolivorans* and *C. fermentans*, and closely related unclassified bacteria (*Clostridium* sp. W14A, *Caproiciproducens* sp. NJN-50, and *Clostridium* sp. KNHs216), we would classify strain 7D4C2 and *C. fermentans* into the same genus of *Caproiciproducens* with *C. galactitolivorans*. The comparable phenotype and similar chain-elongation metabolism between strain 7D4C2, C. *fermentans*, and *C. galactitolivorans* also support that these bacteria belong to the same genus. Thus, we name our isolate *Caproiciproducen*s *fermentans* 7D4C2, which is the same species as *Clostridium* sp. W14A and *C. fermentans*. The rBOX genes of these *Caproiciproducens* species are highly similar and relatively distant to the genes of other chain-elongating bacteria. The 6 rBOX genes in the *Caproiciproducens* spp. are located next to each other, forming a gene cluster. This rBOX cluster is also present in bacteria that do not chain elongate such as *A. senegalensis* and *Clostridium* spp.. The close similarity of the rBOX genes of strain 7D4C2 with these bacteria requires further investigation to understand what defines a chain elongator.

## Materials and Methods

### Isolation of strain 7D4C2

Rumen fluid (from a young sheep) and thermophilic anaerobic sludge, which was collected at the Western Lake Superior Sanitary District in 2011 (Duluth, MN, USA), were used to inoculate a bioreactor converting pretreated cellulosic hydrolysate into *n*-butyrate (Agler et al., 2012b).

Mixed liquor from this bioreactor was used to start a chain-elongation study with ethanol beer (Agler et al., 2012a; Ge et al., 2015). After 5 years of chain elongation with ethanol beer, the mixed liquor was used to inoculate three chain-elongating bioreactors producing *n*-caproate and *n*-caprylate from ethanol and acetate (Spirito *et al*., unpublished data). We used a cryogenic sample from one of these reactors to isolate bacteria *via* soft agar serial dilutions, as indicated in **Additional File 1: Figure S1**. For this, 10 mL of sterile and reduced supplemented basal medium (s-basal medium) (**Additional File 1: Table S5**), containing 0.6 % w/v Bacto Agar (Becton Dickinson, Sparks, MD, USA), were dispensed in 15-mL test tubes that were capped with butyl rubber stoppers and screw caps. After 1 – 2 weeks of incubation at 30°C and a pH of 5.2 ± 0.1, we picked single colonies in an anaerobic glove box (MBraun, Garching, Germany). We cultured the selected colonies in 10 mL of supplemented basal medium with ethanol (Sigma-Aldrich, Steinheim, Germany) and/or fructose (Carl Roth, Karlsruhe, Germany) as substrates in 50-mL serum bottles. After 1 – 2 weeks of cultivation (when the serum bottles were turbid), we measured *n*-caproate and H_2_ production and substrate consumption. The purity of cultures that produced *n*-caproate was examined through scanning electron and/or light microscopy and Sanger sequencing. The isolate that showed 100% purity is referred to as strain 7D4C2.

### Cultivation of strain 7D4C2

We evaluated the chain-elongating metabolism of strain 7D4C2 with different electron acceptors, as well as its *n*-caproate and lactate production at different pH values and temperatures. For these, we grew strain 7D4C2 in 50-mL serum bottles with 10 mL of s-basal medium buffered with 93.18 ± 6.85 mM MES (Carl Roth, Karlsruhe, Germany) (**Additional File 1: Table S5**). For the electron-acceptor experiment (30°C, pH 5.5 ± 0.02), we used 24.4 ± 1.7 mM fructose (146.4 ± 10.3 mmol C L^−1^) and the following carboxylates at a concentration of 108.2 ± 8.0 mmol C L^−1^: Na-acetate (VWR, Solon, OH, USA), Na-butyrate, propionic acid (Merck, Darmstadt, Germany), *n*-valeric acid (Merck, Darmstadt, Germany) and *n*-caproic acid (Carl Roth, Karlsruhe, Germany). This experiment was performed in triplicates. In the experiments at different pH values and temperatures, the primary substrates were 24.7 ± 0.5 mM fructose (148.2 ± 3.2 mmol C L^−1^) and 18.7 ± 1.0 mM Na-butyrate (112.2 ± 6.3 mmol C L^−1^) (ThermoFisher, Kandel, Germany). For the pH experiment, we grew strain 7D4C2 at 30°C in a pH range from 4.5 to 10. The initial pH value was adjusted with 2 N sodium hydroxide (Sigma Aldrich, Steinheim, Germany). For the temperature test, we grew strain 7D4C2 at various temperatures (*i*.*e*., 22.5°C, 27°C, 30°C, 37°C, 42°C, 50°C) at the previously determined optimum pH value (*i*.*e*., pH of 6.0). These experiments were performed in duplicates.

### Extraction of *n*-caproate with mineral oil and 3% (w/v) TOPO

To assess whether the bacterium could produce more *n*-caproate without the inhibition of the undissociated acid, we continuously extracted the MCC using an extraction solvent. The extraction solvent consisted of 30 g/L of tri-n-octylphosphine oxide (TOPO, Acros Organics, Geel, Belgium) in mineral oil (Sigma Aldrich, Steinheim, Germany) (Kucek et al., 2016a). For this experiment, we grew strain 7D4C2 in 50-mL serum bottles containing 10 mL of s-basal medium (314.1 ± 2.1 mmol C L^−1^ fructose, 101.3 ± 3.2 mmol C L^−1^ Na-butyrate, pH 5.2) (**Additional File 1: Table S5**). We added 10 mL of UV light-sterilized extraction solvent after three days of growth, when the *n*-caproate concentration was increasing, to prevent the initial loss of substrate (*i*.*e*., *n*-butyrate) into the extractant. The solvent preferentially extracts hydrophobic molecules, resulting in extraction efficiencies of 83–93% for MCCs and 5–31% for SCCs (Agler et al., 2012b). Because *n*-caproate is more hydrophobic than *n*-butyrate when *n*-caproate is present, it is the main carboxylate extracted. The control serum bottles did not include an extraction solvent. Along with the addition of extractant, we added ∼30 mM more fructose into all serum bottles to promote *n*-caproate production. We calculated the concentration of undissociated acid using the Henderson-Hasselback equation (Harroff et al., 2017). We took liquid samples (0.6 mL) from the culture and solvent phases. We washed the solvent samples five times with an equal amount of 0.3 M sodium borate (Acros Organics, Geel, Belgium) (pH = 9) to back-extract the carboxylic acids. The aqueous phase (*i*.*e*., boric acid with the extracted carboxylates) of each wash was analyzed as indicated below. The concentrations from each washing were summed to estimate the carboxylate production/consumption per data point. We tested these experiments in triplicates at 30°C.

### Comparison among strain 7D4C2, *C. galactitolivorans*, and [*C*.] *leptum*

*C. galactitolivorans* BS-1 was acquired from the Japan Collection of Microorganisms RIKEN and [*C*.] *leptum* VPI T7-24-1 from the German Collection of Microorganisms and Cell Cultures (DSMZ). The sugar consumption of strain 7D4C2, C. *galactitolivorans*, and *C. leptum* was compared in 50-mL serum bottles incubated at 37°C and a pH of 7.0. Since *C. leptum* did not grow in the supplemented basal medium in which we grew strain 7D4C2 (**Additional File 1: Table S5**), nor in the optimized medium for *C. galactitolivorans* (Jeon et al., 2013), the three bacteria were grown in 10 mL of DSMZ medium 107c with glucose as the primary substrate. We tested these experiments in triplicates.

### Analysis of sugars, carboxylates, and H_2_

We quantified sugars and carboxylates (the total of the dissociated and undissociated forms) throughout the culturing period *via* high-performance liquid chromatography (HPLC), as described in (Klask et al., 2020). For the sample preparation, 0.6 mL culture were centrifuged at 13,350 rpm for 6 min in a Benchtop centrifuge (5424 Eppendorf, Hamburg, Germany). The supernatant was filtered through a 0.22-µm polyvinylidene fluoride syringe filter (Carl Roth, Karlsruhe, Germany) and stored alongside the biomass pellets at −20°C until analyzed. Only the acetate, *n*-butyrate, and *n*-caproate concentrations from the pH experiment were analyzed with an Agilent 7890B Gas Chromatograph (Agilent Technologies, Inc., Santa Clara, CA, USA), equipped with a capillary column (DB-Fatwax UI 30 m x 0.25 m; Agilent Technologies) and an FID detector with a ramp temperature program (initial temperature of 80°C for 0.5 min, then 20°C per min up to 180°C, and final temperature of 180°C for 1 min). The injection and detector temperatures were 250°C and 275°C, respectively. Samples were prepared as for HPLC with the addition of an internal standard (Ethyl-butyric acid) and acidification (to pH 2) with 50% formic acid.

To assess H_2_ production, we collected 250-µL gas samples with a 500-µL syringe (Hamilton, Giarmata, Romania). We injected 200 µL in a gas chromatograph (SRI 570 8610C, SRI Instruments, Las Vegas, NV, USA) with the characteristics described in (Ruaud et al., 2020). We used the ideal gas equation to calculate the moles of H_2_ produced per culture volume. For this, we measured the gas pressure in the serum bottles with a digital pressure gauge (Cole Parmer, Vernon Hills, IL, USA). We measured the cell density (OD_600_) with a NanoPhotometer NP80 at 600 nm with a path length of 0.67 mm (Implen, Westlake Village, CA, USA).

### Microscopy and morphology characterization

To image the isolate *via* light microscopy, we centrifuged a 0.5-mL sample of culture in the exponential phase at 7000 rpm for 5 min in a Benchtop centrifuge (5424 Eppendorf, Hamburg, Germany). We washed the pelleted biomass 1 – 2 times and resuspended it with 50 µL 1x PBS from which we fixed 2 µL on solidified agarose (VWR, Solon, OH, USA) (1% w/v). To image the isolate *via* Scanning Electron Microscopy (SEM), we pelleted 6 mL of culture for 3 min at 7000 rpm (Benchtop centrifuge 5424 Eppendorf, Hamburg, Germany) inside a glove-box (MBraun, Garching, Germany). We washed the pellet five times with 500 µL of 1x PBS. After the last washing step, we resuspended the pellet with 450 µL of 1x PBS and added 50 µL of 25% (v/v) glutaraldehyde for fixation. Samples were incubated at room temperature for 2 h, and then handed over to the SEM center at the Max-Planck Institute for Developmental Biology (Tübingen, Germany) for further processing and imaging, as detailed in Ruaud et al. (2020). For Gram staining, we used the Gram stain for films kit (Sigma-Aldrich, Steinheim, Germany), as described in the manufacturer’s protocol.

### DNA extraction and 16S rRNA gene sequence phylogenetic analysis

We extracted DNA from the biomass pellets stored at −20°C using a NucleoSpin*®* Microbial DNA Kit (Macherey-Nagel, Düren, Deutschland), according to the manufacturer’s protocol. The 16S rRNA gene was amplified from genomic DNA using the universal primers sets 27F/1391R and 27F/1525R. The PCR product was purified with DNA Clean Concentrator-5 (Zymo Research, Irvine, CA, USA). Universal primers 27F, 342F, 515F, 926F, and 926R and the designed primer 1492-capro-R (CTACCTTGTTACGACTTCACC) were used to sequence the whole 16S rRNA gene *via* Sanger sequencing. We designed primer 1492-capro-R using the 16S rRNA gene sequence of *C. galactitolivorans* (NCBI <u>FJ805840</u>) as reference. PCR products were sent for sequencing to the Genome Center at the MPI for Developmental Biology (Tübingen, Germany). We used Geneious Prime*®* 2019.1.3 (http://www.geneious.com) to trim and align the DNA sequences, using the global Geneious alignment tool at a 93% similarity with gap open and gap extension penalties of 8 and 2, respectively, and 15 refinement iterations. We compared the assembled 16S rRNA gene sequence to the four sequences extracted from the genome using the Basic Local Alignment Search Tool (BLAST) from the National Center for Biotechnology Information (NCBI) (https://blast.ncbi.nlm.nih.gov/Blast.cgi). We used the most similar sequence (1517 bp) to the Sanger assembly (99.46%) to construct a phylogenetic tree of strain 7D4C2 and its closest relatives. For this, we aligned the 16S rRNA gene sequence to sequences in the Standard nucleotide collection (nr/nt) database using the NCBI BLAST. We constructed the phylogenetic tree using the Single-Genes-Tree tool (http://ggdc.dsmz.de/).

Pairwise sequence similarities between the 16S rRNA gene and closest relatives were calculated using the method recommended by (Meier-Kolthoff et al., 2013b) for the 16S rRNA gene sequence available *via* the Genome to Genome Distance Calculator (GGDC) web server (Meier-Kolthoff et al., 2013a) accessible at http://ggdc.dsmz.de/. Phylogenies were inferred by the GGDC web server (Meier-Kolthoff et al., 2013a), using the DSMZ phylogenomics pipeline (Meier-Kolthoff et al., 2014), which was adapted to single genes. A multiple-sequence alignment was created with MUSCLE (Edgar 2004). Maximum likelihood (ML) and maximum parsimony (MP) trees were inferred from the alignment with RAxML (Stamatakis, 2014) and TNT(Goloboff et al., 2008), respectively. For ML, rapid bootstrapping in conjunction with the autoMRE bootstrapping criterion (Pattengale et al., 2010) and subsequent search for the best tree was used. For MP, 1000 bootstrapping replicates were used in conjunction with tree-bisection-and-reconnection branch swapping and ten random sequence addition replicates. The sequences were checked for a compositional bias using the X^2^ test as implemented in PAUP* (Swofford, 2002).

### Genome sequencing, assembly, alignment, and annotations

The DNA was extracted using a NucleoSpin® Microbial DNA Kit (Macherey-Nagel, Düren, Deutschland), according to the manufacturer’s protocol. The DNA library was prepared using a Rapid barcoding kit (SQK-RBK004, Oxford Nanopore Technologies Ltd., Oxford Science Park, UK). The DNA was sequenced using a MinION sequencer (Oxford Nanopore Technologies Ltd., Oxford Science Park, UK) with a single R9.4.1 flow cell. The basecalling was performed with guppy (v 3.6.0) in high accuracy mode. The basecalled reads were assembled using Unicycler (Wick et al., 2017) (v 0.4.8) with three rounds of Racon (Vaser et al., 2017) (v 1.4.10) polishing, and one round of medaka (v 1.0.1 - https://github.com/nanoporetech/medaka) correction in r941_min_high_g360 mode. The corrected assembly resulted in a single, circular, closed chromosome. The quality of the assembly (contamination and completeness) was assessed using CheckM in lineage_wf mode (Parks et al., 2015). We annotated the assembled chromosome using PGAP (Tatusova et al., 2016) (v 2020-03-30.build4489). We obtained 3914 genes in total. The products of the 722 of the 3633 (19.9%) CDS were annotated as “hypothetical protein”. We aligned the predicted CDSs against EggNOG 5.0 (Huerta-Cepas et al., 2019) database, using eggnog-mapper (Huerta-Cepas et al., 2017) (v 2.0.1) with DIAMOND as the choice of the aligner, and assigned a COG annotation to 3338 of them (91.8%).

### Taxonomic placement

To assign taxonomy, we extracted the identified 16S rRNA gene sequences and aligned them against the National Center for Biotechnology Information nucleotide database (NCBI-nt). We aligned the whole chromosome against NCBI-nt using minimap2 (Li, 2018) (in asm20 mode) and against NCBI-nr (protein database) using DIAMOND (Buchfink et al., 2015) (with the --long-reads parameter), and assigned taxonomy to it using MEGAN-LR (Huson et al., 2018) (with parameters --lcaCoveragePercent 51 and --longReads). We also used GTDB-Tk (Chaumeil et al., 2020) to classify the genome using the Genome Taxonomy Database (Parks et al., 2018).

All methods agreed on assigning strain 7D4C2 to the unclassified organism *Clostridium* sp. W14A. To further explore the taxonomy of strain 7D4C2, we calculated its average nucleotide identity (ANI) using JSpeciesWS (Richter et al., 2016) to all genomes available for the Clostridiales class in GenBank (8662 genomes, accessed on 07/11/2019). We chose the 13 most similar classified microbes for further analysis and used *C. kluyveri* as an outgroup. Next, we compared the percentage of conversed proteins (POCP) as proposed in (Qin et al., 2014), and the genome relatedness index as proposed in (Barco et al., 2020).

### Phylogenetic analysis and synteny of the genes in the rBOX cluster

We aligned the genes from strain 7D4C2 that are known to be responsible for chain elongation (*i*.*e*., *thl, hbd, crt, acdh*, and *etf-*α and *-*β) against the protein sets of closely related microbes, using DIAMOND (Buchfink et al., 2015) (more-sensitive setting) in BLASTP mode. We obtained the homologs of these proteins in the genomes of bacteria closely related to strain 7D4C2 by filtering DIAMOND hits that cover more than 90% of the query and have more than 45% of positives in the alignment. Because some bacteria had several genes coding for rBOX proteins, for our phylogenetic analyses we focused on the genes that formed a cluster or on those most similar to the genes considered from other bacteria. We computed multiple sequence alignments of the rBOX homologs using MUSCLE (Edgar, 2004) and phylogenetic trees using RAxML (Stamatakis, 2014) with 1000 rounds of bootstrapping (PROTGAMMAAUTO model, parsimony seed set to 12345). We also generated a consensus tree using SplitsTree 5 (v 5.0.0_alpha, with Consensus=Greedy option) (Huson, 1998) of all of the 17 taxa and 6 gene trees. We traced back the genomic coordinates of the rBOX homologs from their annotations on NCBI RefSeq, and used this information to check for synteny and their organization in the genomes manually.

## Additional Files

### Additional File 1

Figure S1: Summary of the isolation process of strain 7D4C2.

Figure S2: Gram staining of strain 7D4C2 and controls for negative and positive staining. Figures S3: OD_600_ and H_2_ production throughout the culturing period for strain 7D4C2 at different pH values and temperatures.

Figures S4: Fructose, *n*-butyrate, and products concentrations throughout the culturing period for strain 7D4C2 at different pH values and temperatures.

Figure S5: OD_600_ and H_2_ production throughout the culturing period for strain 7D4C2 with and without extraction solvent.

Figure S6: Phylogeny between strain 7D4C2 and its closest relatives based on the 16S rRNA gene sequence.

Figure S7: Comparison of glucose fermentation by strain 7D4C2, *C. galactitolivorans* and *C. leptum*.

Figure S8: Substrate consumption by strain 7D4C2 according to the AN MicroPlate™ from Biolog (Hayward, CA.)

Figure S9: Phylogenetic trees of the rBOX genes in strain 7D4C2, closest relatives, and known chain-elongating bacteria.

Table S1: Maximum OD_600_, final electron donor and acceptor and product concentrations, and *n*-caproate specificity in cultures of strain 7D4C2 with different electron acceptors.

Table S2: Maximum OD_600_ values, final concentration of products, and specificities of lactate and *n*-caproate in cultures of strain 7D4C2 grown at different pH values and temperatures. Table S3. Average nucleotide identity (ANI) and alignment fraction (AF) values between strain 7D4C2 and most similar strains.

Table S4: Comparison of carbohydrates oxidized by strain 7D4C2, *C. galactitolivorans*, and *C. fermentans*.

Table S5: Composition of the supplemented basal medium.

### Additional File 2

Location and percent identity of all rBOX genes in strain 7D4C2, closely related bacteria, bacteria with similar rBOX genes, and known chain-elongating bacteria. In green: rBOX genes used in the phylogenetic analyses.

## List of abbreviations

MCC: medium-chain carboxylate (comprising both the dissociated and undissociated forms)
SCC: short-chain carboxylate (comprising both the dissociated and undissociated forms)
rBOX: reverse β-oxidation
Thl: thiolase
HBD: 3-hydroxybutyryl-CoA dehydrogenase
Crt: crotonyl-CoA
ACDH: acyl-CoA dehydrogenase
ETF: electron transport flavoprotein
ANI: average nucleotide identity
POCP: percentage of conserved proteins
AF: aligned fraction
OD_600_: optical density measured at 600 nm.

## Availability of data and materials

Strain 7D4C2 was deposited in the German Collection of Microorganisms and Cell Cultures (DSMZ) under the accession number DSM 110548. The datasets generated and analyzed during the present study are included in this published article and are available from LA and SEE on request. The assembled 16S rRNA sequences and the whole-genome of the isolate are available online (https://www.ncbi.nlm.nih.gov) under the accession numbers NCBI <u>MT056029</u> and Project ID PRJNA615378, respectively. Raw sequencing MinION data are available online (https://www.ncbi.nlm.nih.gov) under the project ID.

## Authors contributions

LTA conceived the project, and SEE designed and guided the study. MT and SEE performed the lab experiments. CB performed the bioinformatics analyses. MT, CB, and SEE analyzed the data. MT, CB, and SEE prepared the figures and tables. SEE, LTA, CB, and MT drafted the manuscript. BYJ and IB performed the genome sequencing, and RBHW advised on the sequencing tools. LTA and DHH provided guidance. All authors edited the manuscript and approved the final manuscript.

## Acknowledgments

Authors are thankful to Ursula Schach for the valuable help acquiring *C. galactitolivorans* and dealing with the deposit agreements with the DSMZ and ATCC collection banks.

## References

Agler, M.T., Spirito, C.M., Usack, J.G., Werner, J.J., Angenent, L.T. 2012a. Chain elongation with reactor microbiomes: upgrading dilute ethanol to medium-chain carboxylates. Energy Environ. Sci., 5(8), 8189.

Agler, M.T., Spirito, C.M., Usack, J.G., Werner, J.J., Angenent, L.T. 2014. Development of a highly specific and productive process for n-caproic acid production: applying lessons from methanogenic microbiomes. Water Sci. Technol., 69(1), 62–68.

Agler, M.T., Werner, J.J., Iten, L.B., Dekker, A., Cotta, M.A., Dien, B.S., Angenent, L.T. 2012b. Shaping reactor microbiomes to produce the fuel precursor n-butyrate from pretreated cellulosic hydrolysates. Environ. Sci. Technol., 46(18), 10229–10238.

Angenent, L.T., Richter, H., Buckel, W., Spirito, C.M., Steinbusch, K.J.J., Plugge, C.M., Strik, D.P.B.T.B., Grootscholten, T.I.M., Buisman, C.J.N., Hamelers, H.V.M. 2016. Chain elongation with reactor microbiomes: Open-culture biotechnology to produce biochemicals. Environ. Sci. Technol., 50(6), 2796–2810.

Barco, R., Garrity, G., Scott, J., Amend, J., Nealson, K., Emerson, D. 2020. A genus definition for bacteria and archaea based on a standard genome relatedness index. mBio, 11(1).

Buchfink, B., Xie, C., Huson, D.H. 2015. Fast and sensitive protein alignment using DIAMOND. Nat. methods, 12(1), 59.

Chaumeil, P.-A., Mussig, A.J., Hugenholtz, P., Parks, D.H. 2020. GTDB-Tk: a toolkit to classify genomes with the Genome Taxonomy Database, Oxford University Press.

Contreras-Dávila, C.A., Carrión, V.J., Vonk, V.R., Buisman, C.N.J., Strik, D.P.B.T.B. 2020. Consecutive lactate formation and chain elongation to reduce exogenous chemicals input in repeated-batch food waste fermentation. Water Res., 169, 115215.

Desbois, A.P. 2012. Potential applications of antimicrobial fatty acids in medicine, agriculture and other industries. Recent Pat. Antiinfect. Drug Discov., 7(2), 111–122.

Duber, A., Jaroszynski, L., Zagrodnik, R., Chwialkowska, J., Juzwa, W., Ciesielski, S., Oleskowicz-Popiel, P. 2018. Exploiting the real wastewater potential for resource recovery – *n*-caproate production from acid whey. Green Chem., 20(16), 3790–3803.

Edgar, R.C. 2004. MUSCLE: multiple sequence alignment with high accuracy and high throughput. Nucleic Acids Res., 32(5), 1792–1797.

Felicity A. Roddick, Britx, M.L. 1997. Production of hexanoic acid by free and immobilised cells of *Megasphaera elsdenii*: Influence of *in-situ* product removal using ion exchange resin. J. Chem. Tech. Biotechnol, 69, 383–391.

Flaiz, M., Baur, T., Brahner, S., Poehlein, A., Daniel, R., Bengelsdorf, F.R. 2020. *Caproicibacter fermentans* gen. nov., sp. nov., a new caproate-producing bacterium and emended description of the genus Caproiciproducens. Int. J. Syst. Evol. Microbiol.

Ge, S., Usack, J.G., Spirito, C.M., Angenent, L.T. 2015. Long-term *n*-caproic acid production from yeast-fermentationbBeer in an anaerobic bioreactor with continuous product extraction. Environ. Sci. Technol., 49(13), 8012–8021.

Goloboff, P.A., Farris, J.S., Nixon, K.C. 2008. TNT, a free program for phylogenetic analysis. Cladistics, 24(5), 774–786.

Harroff, L.A., Liotta, J.L., Bowman, D.D., Angenent, L.T. 2017. Inactivation of ascaris eggs in human fecal material through *in situ* production of carboxylic acids. Environ. Sci. Technol., 51(17), 9729–9738.

Harvey, B.G., Meylemans, H.A. 2014. 1-Hexene: a renewable C6 platform for full-performance jet and diesel fuels. Green Chem., 16(2), 770–776.

Huerta-Cepas, J., Forslund, K., Coelho, L.P., Szklarczyk, D., Jensen, L.J., Von Mering, C., Bork, P. 2017. Fast genome-wide functional annotation through orthology assignment by eggNOG-mapper. Mol. Biol. Evol., 34(8), 2115–2122.

Huerta-Cepas, J., Szklarczyk, D., Heller, D., Hernández-Plaza, A., Forslund, S.K., Cook, H., Mende, D.R., Letunic, I., Rattei, T., Jensen, L.J. 2019. eggNOG 5.0: a hierarchical, functionally and phylogenetically annotated orthology resource based on 5090 organisms and 2502 viruses. Nucleic Acids Res., 47(D1), D309–D314.

Huson, D.H. 1998. SplitsTree: analyzing and visualizing evolutionary data. Bioinformatics (Oxford, England), 14(1), 68–73.

Huson, D.H., Albrecht, B., Bağcı, C., Bessarab, I., Gorska, A., Jolic, D., Williams, R.B. 2018. MEGAN-LR: new algorithms allow accurate binning and easy interactive exploration of metagenomic long reads and contigs. Biol. Direct, 13(1), 6.

Jeon, B.S., Choi, O., Um, Y., Sang, B.-I. 2016. Production of medium-chain carboxylic acids by *Megasphaera* sp. MH with supplemental electron acceptors. Biotechnol. Biofuels, 9, 129.

Jeon, B.S., Kim, B.-C., Um, Y., Sang, B.-I. 2010. Production of hexanoic acid from D-galactitol by a newly isolated *Clostridium* sp. BS-1. Appl. Microbiol. Biotechnol., 88(5), 1161–1167.

Jeon, B.S., Moon, C., Kim, B.-C., Kim, H., Um, Y., Sang, B.-I. 2013. *In situ* extractive fermentation for the production of hexanoic acid from galactitol by *Clostridium* sp. BS-1. Enzyme Microb. Tech., 53(3), 143–151.

Kenealy, W.R., Cao, Y., Weimer, P.J. 1995. Production of caproic acid by cocultures of ruminal cellulolytic bacteria and *Clostridium kluyveri* grown on cellulose and ethanol. Appl. Microbiol. Biotechnol., 44(3), 507–513.

Kim, B.-C., Seung Jeon, B., Kim, S., Kim, H., Um, Y., Sang, B.-I. 2015. *Caproiciproducens galactitolivorans* gen. nov., sp. nov., a bacterium capable of producing caproic acid from galactitol, isolated from a wastewater treatment plant. Int. J. Syst. Evol. Microbiol., 65(12), 4902–4908.

Klask, C.-M., Kliem-Kuster, N., Molitor, B., Angenent, L.T. 2020. Nitrate feed improves growth and ethanol production of *Clostridium ljungdahlii* with CO_2_ and H_2_, but results in stochastic inhibition events. Front. Microbiol., 11, 724.

Kucek, L.A., Nguyen, M., Angenent, L.T. 2016a. xsConversion of L-lactate into *n*-caproate by a continuously fed reactor microbiome. Water Res., 93, 163–171.

Kucek, L.A., Spirito, C.M., Angenent, L.T. 2016b. High *n*-caprylate productivities and specificities from dilute ethanol and acetate: chain elongation with microbiomes to upgrade products from syngas fermentation. Energy Environ. Sci., 9(11), 3482–3494.

Lanjekar, V.B., Marathe, N.P., Ramana, V.V., Shouche, Y.S., Ranade, D.R. 2014. *Megasphaera indica* sp. nov., an obligate anaerobic bacteria isolated from human faeces. Int. J. Syst. Evol. Microbiol., 64(Pt 7), 2250–2256.

Levy, P.F., Sanderson, J.E., Kispert, R.G., Wise, D.L. 1981. Biorefining of biomass to liquid fuels and organic chemicals. Enzyme Microb. Techn., 3(3), 207–215.

Li, H. 2018. Minimap2: pairwise alignment for nucleotide sequences. Bioinformatics, 34(18), 3094–3100.

Lindley, N., Loubiere, P., Pacaud, S., Mariotto, C., Goma, G. 1987. Novel products of the acidogenic fermentation of methanol during growth of *Eubacterium limosum* in the presence of high concentrations of organic acids. Microbiol., 133(12), 3557–3563.

Lino, T., Mori, K., Tanaka, K., Suzuki, K.-i., Harayama, S. 2007. *Oscillibacter valericigenes* gen. nov., sp. nov., a valerate-producing anaerobic bacterium isolated from the alimentary canal of a Japanese corbicula clam. I nt. J. Syst. Evol. Microbiol., 57(8), 1840–1845.

Marounek, M., Fliegrova, K., Bartos, S. 1989. Metabolism and some characteristics of ruminal strains of *Megasphaera elsdenii*. Appl. Environ. Microbiol., 55(6), 1570–1573.

Meier-Kolthoff, J.P., Auch, A.F., Klenk, H.-P., Göker, M. 2013a. Genome sequence-based species delimitation with confidence intervals and improved distance functions. BMC Bioinformatics, 14, 60.

Meier-Kolthoff, J.P., Göker, M., Spröer, C., Klenk, H.-P. 2013b. When should a DDH experiment be mandatory in microbial taxonomy? Arch. Microbiol., 195(6), 413–418.

Meier-Kolthoff, J.P., Hahnke, R.L., Petersen, J., Scheuner, C., Michael, V., Fiebig, A., Rohde, C., Rohde, M., Fartmann, B., Goodwin, L.A., Chertkov, O., Reddy, T.B.K., Pati, A., Ivanova, N.N., Markowitz, V., Kyrpides, N.C., Woyke, T., Göker, M., Klenk, H.-P. 2014. Complete genome sequence of DSM 30083T, the type strain (U5/41T) of *Escherichia coli*, and a proposal for delineating subspecies in microbial taxonomy. Stand. Genomic Sci., 9(1), 2.

Park, S., Yasin, M., Jeong, J., Cha, M., Kang, H., Jang, N., Choi, I.-G., Chang, I.S. 2017. Acetate-assisted increase of butyrate production by *Eubacterium limosum* KIST612 during carbon monoxide fermentation. Biores. Technol., 245, 560–566.

Parks, D.H., Chuvochina, M., Waite, D.W., Rinke, C., Skarshewski, A., Chaumeil, P.-A., Hugenholtz, P. 2018. A standardized bacterial taxonomy based on genome phylogeny substantially revises the tree of life. Nat. Biotechnol., 36(10), 996–1004.

Parks, D.H., Imelfort, M., Skennerton, C.T., Hugenholtz, P., Tyson, G.W. 2015. CheckM: assessing the quality of microbial genomes recovered from isolates, single cells, and metagenomes. Genome Res., 25(7), 1043–1055.

Pattengale, N.D., Alipour, M., Bininda-Emonds, O.R.P., Moret, B.M.E., Stamatakis, A. 2010. How many bootstrap replicates are necessary? J. Comp. Biol., 17(3), 337–354.

Qin, Q.-L., Xie, B.-B., Zhang, X.-Y., Chen, X.-L., Zhou, B.-C., Zhou, J., Oren, A., Zhang, Y.-Z. 2014. A proposed genus boundary for the prokaryotes based on genomic insights. J. Bacteriol., 196(12), 2210–2215.

Richter, M., Rosselló-Móra, R. 2009. Shifting the genomic gold standard for the prokaryotic species definition. Proc. Natl. Acad. Sci. U. S. A., 106(45), 19126–19131.

Richter, M., Rosselló-Móra, R., Oliver Glöckner, F., Peplies, J. 2016. JSpeciesWS: a web server for prokaryotic species circumscription based on pairwise genome comparison. Bioinformatics, 32(6), 929–931.

Roh, H., Ko, H.-J., Kim, D., Choi, D.G., Park, S., Kim, S., Chang, I.S., Choi, I.-G. 2011. Complete genome sequence of a carbon monoxide-utilizing acetogen, *Eubacterium limosum* KIST612. J. Bacteriol., 193(1), 307–308.

Ruaud, A., Esquivel-Elizondo, S., de la Cuesta-Zuluaga, J., Waters, J.L., Angenent, L.T., Youngblut, N.D., Ley, R.E. 2020. Syntrophy *via* interspecies H_2_ transfer between *Christensenella* and *Methanobrevibacter* underlies their global cooccurrence in the human gut. mBio, 11(1).

Russell, J. 1992. Another explanation for the toxicity of fermentation acids at low pH: anion accumulation versus uncoupling. J. Appl. Bacteriol., 73(5), 363–370.

Spirito, C.M., Marzilli, A.M., Angenent, L.T. 2018. Higher substrate ratios of ethanol to acetate steered chain elongation towards *n*-caprylate in a bioreactor with product extraction. Environ. Sci. Technol., 52(22), 13438–13447.

Spirito, C.M., Richter, H., Rabaey, K., Stams, A.J.M., Angenent, L.T. 2014. Chain elongation in anaerobic reactor microbiomes to recover resources from waste. Curr. Opin. Biotechnol., 27, 115–122.

Stamatakis, A. 2014. RAxML version 8: a tool for phylogenetic analysis and post–analysis of large phylogenies. Bioinformatics, 30(9), 1312–1313.

Swofford, D.L. 2002. PAUP*: Phylogenetic analysis using parsimony (*and other methods). 4.0. B5.

Tatusova, T., DiCuccio, M., Badretdin, A., Chetvernin, V., Nawrocki, E.P., Zaslavsky, L., Lomsadze, A., Pruitt, K.D., Borodovsky, M., Ostell, J. 2016. NCBI prokaryotic genome annotation pipeline. Nucleic Acids Res., 44(14), 6614–6624.

Vaser, R., Sović, I., Nagarajan, N., Šikić, M. 2017. Fast and accurate de novo genome assembly from long uncorrected reads. Genome Res., 27(5), 737–746.

Wang, H., Li, X., Wang, Y., Tao, Y., Lu, S., Zhu, X., Li, D. 2018. Improvement of *n*-caproic acid production with *Ruminococcaceae* bacterium CPB6: selection of electron acceptors and carbon sources and optimization of the culture medium. Microb. Cell Fact., 17(1), 99.

Wick, R.R., Judd, L.M., Gorrie, C.L., Holt, K.E. 2017. Unicycler: resolving bacterial genome assemblies from short and long sequencing reads. PLoS Comput. Biol., 13(6), e1005595.

Xu, J., Guzman, J.J.L., Andersen, S.J., Rabaey, K., Angenent, L.T. 2015. In-line and selective phase separation of medium-chain carboxylic acids using membrane electrolysis. Chem. Commun., 51(31), 6847–6850.

Xu, J., Hao, J., Guzman, J.J.L., Spirito, C.M., Harroff, L.A., Angenent, L.T. 2018. Temperature-phased conversion of acid whey waste into medium-chain carboxylic acids *via* lactic acid: no external e-donor. Joule, 2(2), 280–295.

Yarza, P., Yilmaz, P., Pruesse, E., Glöckner, F.O., Ludwig, W., Schleifer, K.-H., Whitman, W.B., Euzéby, J., Amann, R., Rosselló-Móra, R. 2014. Uniting the classification of cultured and uncultured bacteria and archaea using 16S rRNA gene sequences. Nat. Rev. Microbiol., 12(9), 635–645.

Zhu, X., Zhou, Y., Wang, Y., Wu, T., Li, X., Li, D., Tao, Y. 2017. Production of high-concentration *n*-caproic acid from lactate through fermentation using a newly isolated *Ruminococcaceae* bacterium CPB6. Biotechnol. Biofuels, 10(1).

