## Additional File 1 for "The isolate *Caproiciproducens* sp. 7D4C2 produces *n*-caproate at mildly acidic conditions from hexoses: genome and rBOX comparison with related strains and chain-elongating bacteria"

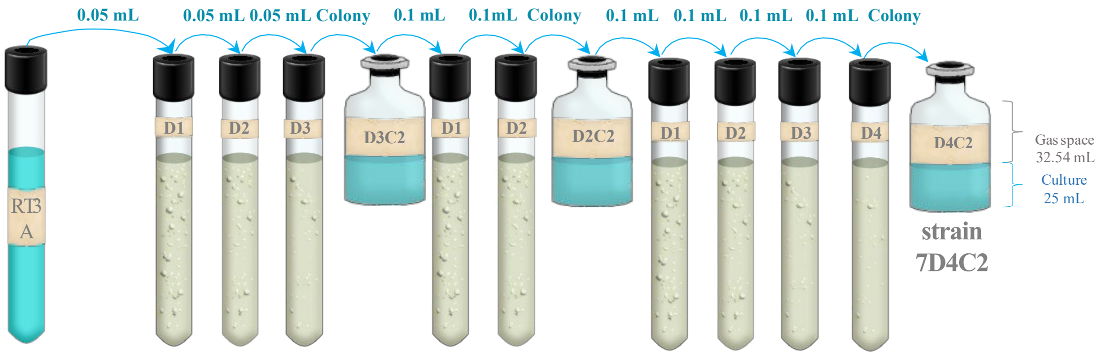


| | Soft agar serial dilutions | Picked colony | Soft agar serial dilutions | Picked colony | Soft agar serial dilutions | Isolate |
|---|---|---|---|---|---|---|
| Ethanol, mM | 50 | 50 | - | - | - | - |
| Acetate, mM | - | - | - | - | 5 | 5 |
| *n*-Butyrate, mM | 57.1 | 57.1 | 15 | 15 | 30 | 30 |
| *n*-Caproate, mM | - | - | - | - | 5 | 5 |
| *n*-Caprylate, mM | 1 | 1 | - | - | - | - |
| Fructose, mM | 55.5 | 55.5 | 30 | 30 | 30 | 30 |
| Yeast extract, % w/v | 0.05 | 0.05 | 0.05 | 0.05 | 0.3 | 0.3 |
| Vitamin sol. 2x, % v/v | 0.5 | 0.5 | 0.5 | 0.5 | 0.2 | 0.2 |
| MES, mM | 91.5 | 91.5 | 91.5 | 91.5 | 91.5 | 91.5 |
| pH | 5.2 | 5.2 | 5.2 | 5.2 | 5.2 | 5.2 |

**Figure S1** Summary of the isolation process of strain 7D4C2. For the isolation, a sample from a chain-elongating reactor (SR1) with in-line product extraction - operated at a pH of 5.5 and a temperature of 30 ºC (Spirito, Angenent et al. unpublished work) - was enriched with ethanol (40 mM), acetate (4 mM), *n*-caproate (4 mM), and *n*-caprylate (4 mM) during 3 transfers (RT3A in the figure). This enriched culture was serially diluted in soft agar with supplemented basal medium (**Table S5**), as indicated in the figure. From the series of dilutions, an *n*-caproate producing colony was further diluted until its purity was confirmed *via* microscopy and Sanger Sequencing. RT3A: Reactor Transfer 3 Replicate A. D: dilution number. C: colony.

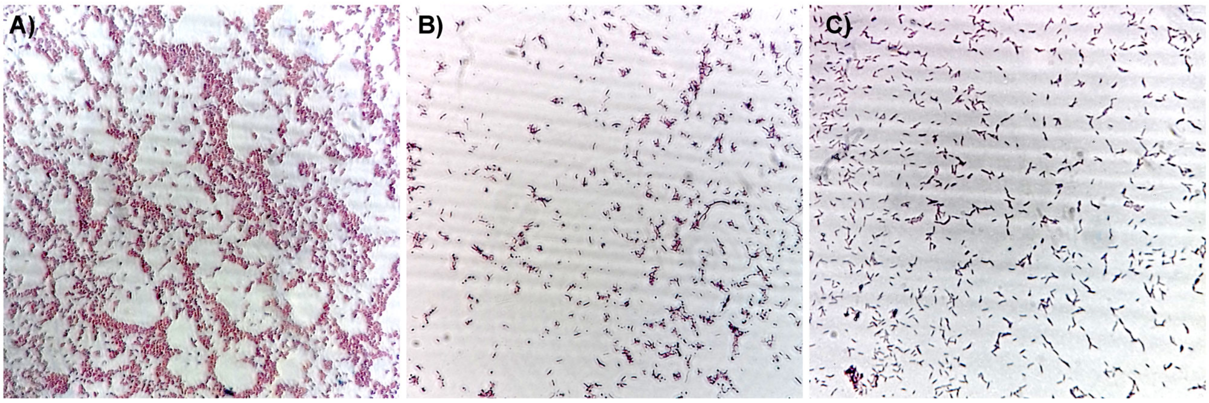


**Figure S2** Gram staining of strain 7D4C2 and controls for negative and positive staining: **A)** *Escherichia colii* NEB® Stable, which is Gram negative); **B)** *Caproiciproducens galactitolivorans* BS-1, which is Gram positive; and **C)** strain 7D4C2, which is found here to be Gram positive.

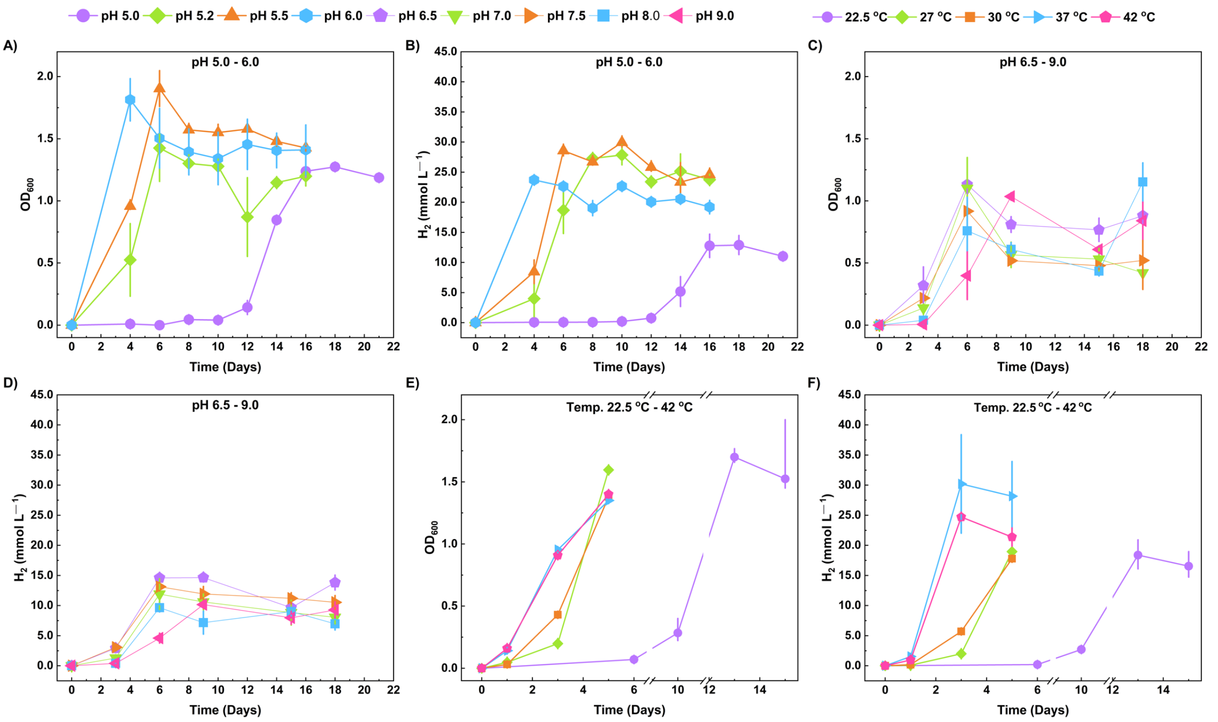


**Figure S3** $OD_{600}$ and $H_2$ production throughout the culturing period for strain 7D4C2 at different pH values and temperatures: **A)** $OD_{600}$ at pH values of 5.0-6.0; **B)** $H_2$ production at pH values of 5.0-6.0; **C)** $OD_{600}$ at pH values of 6.5 to 9.0; **D)** $H_2$ production at pH values of 6.5 to 9.0; **E)** $OD_{600}$ at temperatures of 27 to 42 ºC; and **F)** $H_2$ production at temperatures of 27 to 42ºC. The $OD_{600}$ and $H_2$ production of the test at a pH of 4.5 is not shown due to the extended lag phase of about 20 days. Bars represent minimum and maximum values between duplicate cultures.

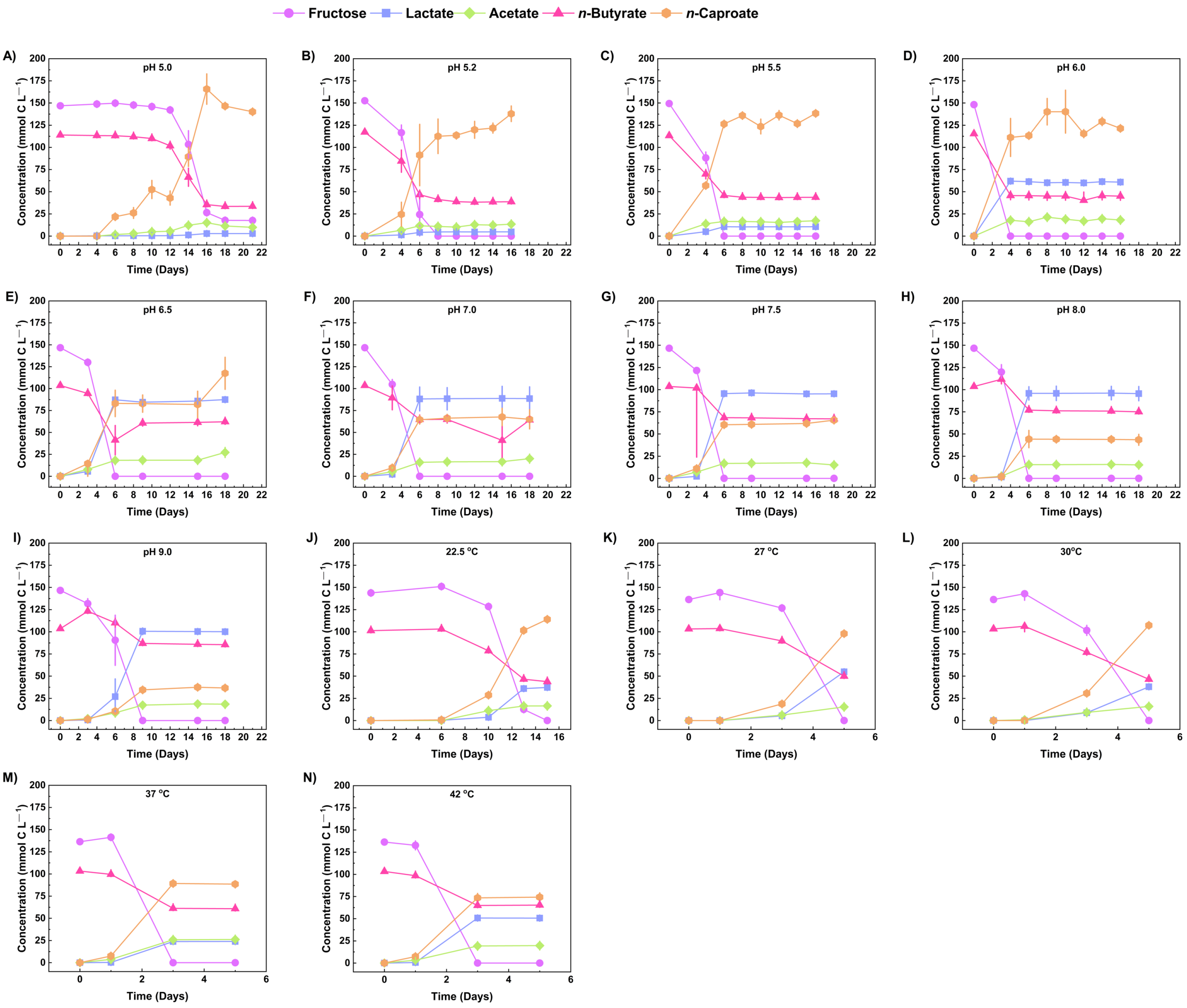


**Figure S4** Fructose, *n*-butyrate, and products concentrations throughout the culturing period for strain 7D4C2 at different pH values and temperatures**: A-I)** graphs at different pH values. The graph at pH of 4.5 is not shown due to the extended lag phase of ~20 days; and **J-N)** graphs at different temperatures. Bars represent minimum and maximum values between duplicate cultures.

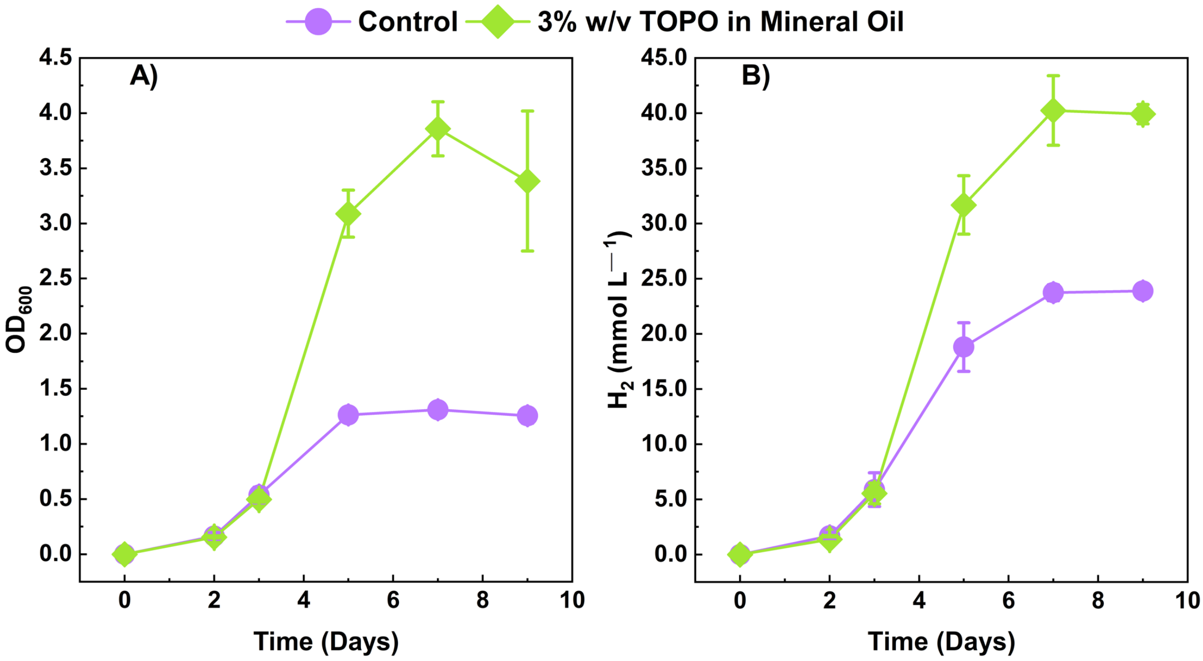


**Figure S5** $OD_{600}$ and $H_2$ production throughout the culturing period for strain 7D4C2 with and without extraction solvent**: A)** $OD_{600}$; and B) $H_2$ production. Error bars represent one standard deviation among triplicate cultures.

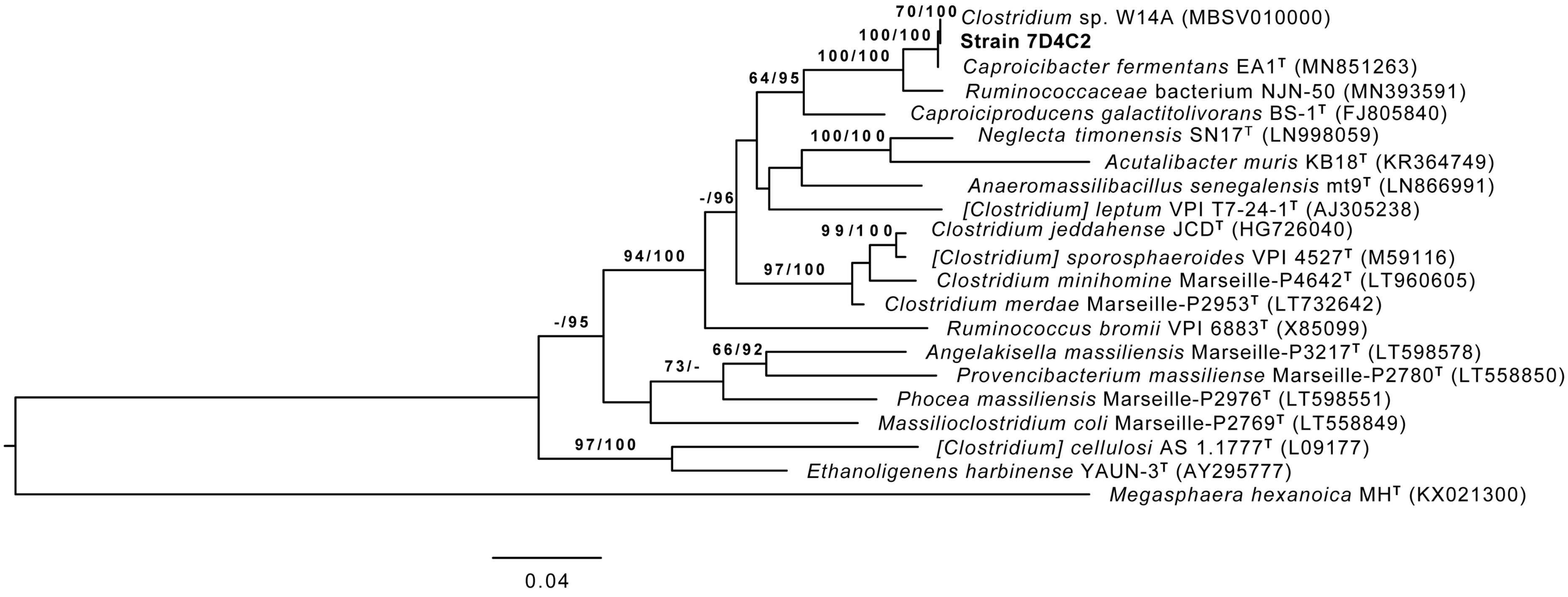


**Figure S6** Phylogeny between strain 7D4C2 and its closest relatives based on the 16S rRNA gene. The branches are scaled in terms of the expected number of substitutions per site. The numbers above the branches are support values when larger than 60% from ML (left) and MP (right) bootstrapping. *Megasphaera hexanoica* MH$^T$ (KX021300) was used as the outgroup. Bar indicates 4.0% sequence divergence. GenBank accession numbers of the 16S rRNA gene sequences are given in parentheses. The tree was built using one of the complete 16S rRNA genes found in the genome (NCBI PRJNA615378, 99.46% similar to the assembly done by Sanger Sequencing, NCBI MT056029).

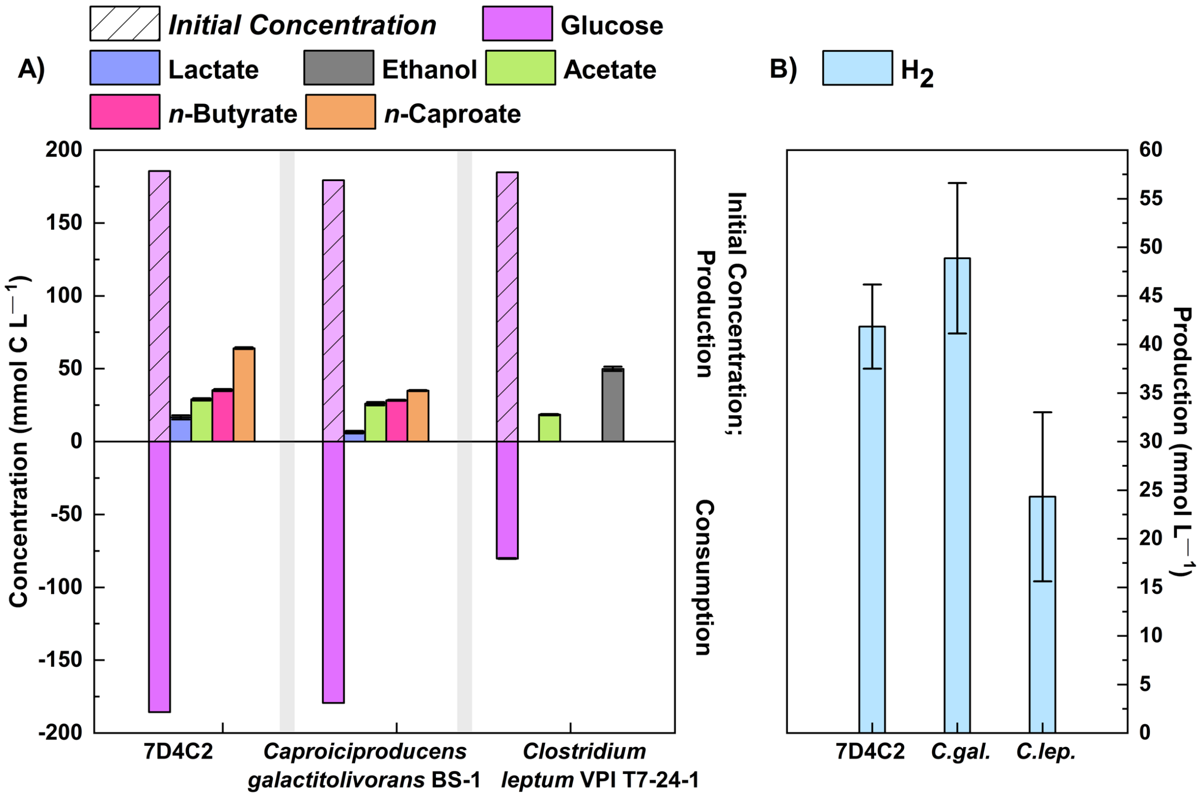


**Figure S7** Comparison of glucose fermentation by strain 7D4C2, *C. galactitolivorans* and *C. leptum* in DSMZ medium 107c (pH 7.0 and 37ºC)**: A)** comparison of final products concentrations and glucose consumption between strains; and B) comparison of final $H_2$ production between strains. Error bars represent one standard deviation among triplicate cultures.

A)

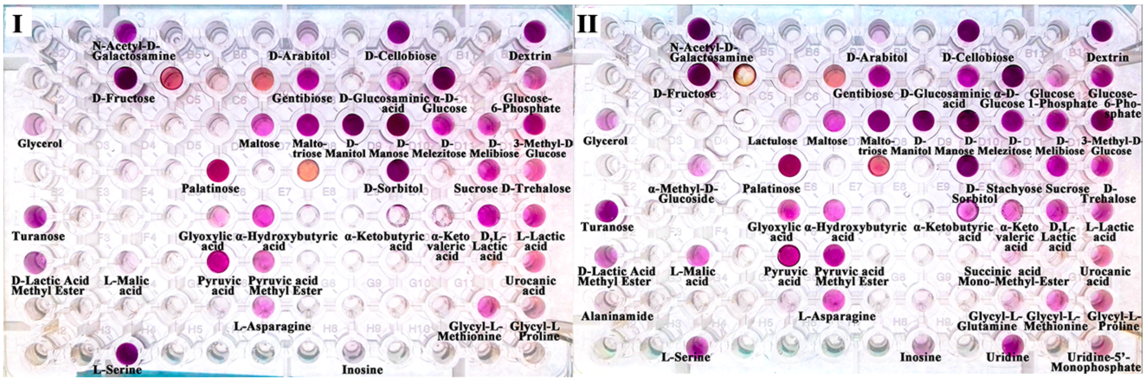


B)

| Substrate | Test I | Test II | Consumption | Substrate | Test I | Test II | Consumption | Substrate | Test I | Test II | Consumption |
|---|---|---|---|---|---|---|---|---|---|---|---|
| Water | 0 | 0 | N | D-Mannose | 1 | 1 | Y | Pyruvic Acid | 1 | 1 | Y |
| N-Acetyl-D- Galactosamine | 0 | 0 | N | D-Melezitose | 1 | 1 | Y | Pyruvic Acid Methyl Ester | 1 | 1 | Y |
| N-Acetyl-D- Glucosamine | 1 | 1 | Y | D-Melibiose | 1 | 1 | Y | D-Saccharic Acid | 0 | 0 | N |
| N-Acetyl-β-D- Mannosamine | 0 | 0 | N | 3-Methyl-D- Glucose | 1 | 1 | Y | Succinamic Acid | 0 | 0 | N |
| Adonitol | 0 | 0 | N | α-Methyl-D- Galactoside | 0 | 0 | N | Succinic Acid | 0 | 0 | N |
| Amygdalin | 0 | 0 | N | β-Methyl-D- Galactoside | 0 | 0 | N | Succinic Acid Mono-Methyl Ester | 0 | 1 | N |
| D-Arabitol | 1 | 1 | Y | α-Methyl-D- Glucoside | 0 | 1 | N | m-Tartaric Acid | 0 | 0 | N |
| Arbutin | 0 | 0 | N | β-Methyl-D- Glucoside | 0 | 0 | N | Urocanic Acid | 1 | 1 | Y |
| D-Cellobiose | 1 | 1 | Y | Palatinose | 1 | 1 | Y | Alaninamide | 0 | 1 | N |
| α-Cyclodextrin | 0 | 0 | N | D-Raffinose | 0 | 0 | N | L-Alanine | 0 | 0 | N |
| β-Cyclodextrin | 0 | 0 | N | L-Rhamnose | 0 | 0 | N | L-Alanyl-L- Glutamine | 0 | 0 | N |
| Dextrin | 1 | 1 | Y | Salicin | 0 | 0 | N | L-Alanyl-L- Histidine | 0 | 0 | N |
| Dulcitol | 0 | 0 | N | D-Sorbitol | 1 | 1 | Y | L-Alanyl-L- Threonine | 0 | 0 | N |
| i-Erythritol | 0 | 0 | N | Stachyose | 0 | 1 | N | L-Asparagine | 1 | 1 | Y |
| D-Fructose | 1 | 1 | Y | Sucrose | 1 | 1 | Y | L-Glutamic Acid | 0 | 0 | N |
| L-Fucose | 0 | 0 | N | D-Trehalose | 1 | 1 | Y | L-Glutamine | 0 | 0 | N |
| D-Galactose | 0 | 0 | N | Turanose | 1 | 1 | Y | Glycyl-L- Aspartic Acid | 0 | 0 | N |
| D-Galacturonic Acid | 0 | 0 | N | Acetic Acid | 0 | 0 | N | Glycyl-L- Glutamine | 0 | 1 | N |
| Gentiobiose | 1 | 1 | Y | Formic Acid | 0 | 0 | N | Glycyl-L- Methionine | 1 | 1 | Y |
| D-Gluconic Acid | 0 | 0 | N | Fumaric Acid | 0 | 0 | N | Glycyl-L- Proline | 1 | 1 | Y |
| D-Glucosaminic Acid | 1 | 1 | Y | Glyoxylic Acid | 1 | 1 | Y | L-Methionine | 0 | 0 | N |
| α-D-Glucose | 1 | 1 | Y | α-Hydroxybutyric Acid | 1 | 1 | Y | L-Phenylalanine | 0 | 0 | N |
| α-D-Glucose-1-Phosphate | 0 | 1 | N | β- Hydroxybutyric Acid | 0 | 0 | N | L-Serine | 1 | 1 | Y |
| D-Glucose-6-Phosphate | 1 | 1 | Y | Itaconic Acid | 0 | 0 | N | L-Threonine | 0 | 0 | N |
| Glycerol | 1 | 1 | Y | α-Ketobutyric Acid | 1 | 1 | Y | L-Valine | 0 | 0 | N |
| D,L-α-Glycerol Phosphate | 0 | 0 | N | α-Ketovaleric Acid | 1 | 1 | Y | L-Valine plus L-Aspartic Acid | 0 | 0 | N |
| m-Inositol | 0 | 0 | N | D,L-Lactic Acid | 1 | 1 | Y | 2'-Deoxy Adenosine | 0 | 0 | N |
| α-D-Lactose | 0 | 0 | N | L-Lactic Acid | 1 | 1 | Y | Inosine | 1 | 1 | Y |
| Lactulose | 0 | 1 | N | D-Lactic Acid Methyl Ester | 1 | 1 | Y | Thymidine | 0 | 0 | N |
| Maltose | 1 | 1 | Y | D-Malic Acid | 0 | 0 | N | Uridine | 0 | 1 | N |
| Maltotriose | 1 | 1 | Y | L-Malic Acid | 1 | 1 | Y | Thymidine-5'-Mono-phosphate | 0 | 0 | N |
| D-Mannitol | 1 | 1 | Y | Propionic Acid | 0 | 0 | N | Uridine-5'- Mono-phosphate | 0 | 1 | N |

**Figure S8** Substrate consumption by strain 7D4C2 according to the AN MicroPlate™ from Biolog (Hayward, CA)**: A)** pictures of the duplicate assays (**I** and **II**), where purple wells indicate oxidation of the substrate; and **B)** summary of results. 1 and 0 indicate positive and negative oxidation, respectively, in each test (**I** and **II**). Only when both tests showed positive oxidation, the substrate was considered to be metabolized (Y).

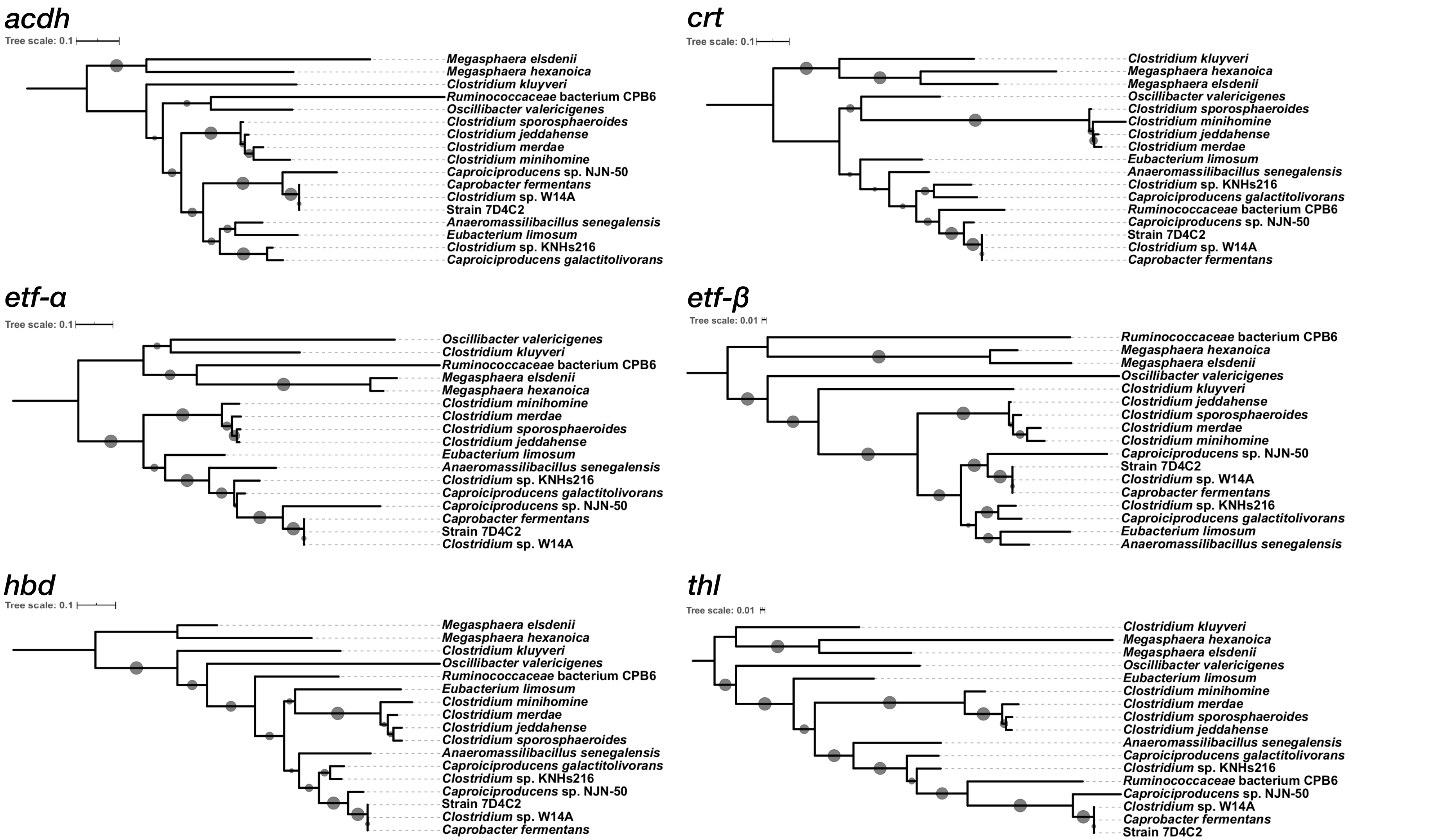


**Figure S9**. Phylogenetic trees of the rBOX genes in strain 7D4C2, closest relatives and known chain-elongating bacteria. The genes of *Clostridium kluyveri* are used as outgroups. The multiple sequence alignments were computed by MUSCLE and the gene trees were inferred by RAxML with 1000 iterations of bootstrapping (values greater than 30% are shown as circles on the edges) and visualized by iTOL.

**Table S1** Maximum $OD_{600}$, final electron donor and acceptor and product concentrations, and *n*-caproate specificity in cultures of strain 7D4C2 with different electron acceptors and for the control with only fructose.

| Electron donor and acceptor | Max $OD_{600}$ | $H_2$ production (mmol $L^{-1}$) | Lactate concentration (mmol C $L^{-1}$) | Acetate concentration (mmol C $L^{-1}$) | Propionate concentration (mmol C $L^{-1}$) | *n*-Butyrate concentration (mmol C $L^{-1}$) | *n*-Valerate concentration (mmol C $L^{-1}$) | *n*-Caproate concentration (mmol C $L^{-1}$) | *n*-Caproate carboxylate specificity (%) |
|---|---|---|---|---|---|---|---|---|---|
| Fruc | 1.59 ± 0.11 | 22.49 ± 2.06 | 25.42 ± 2.69 | 4.56 ± 0.64 | ND | 6.93 ± 0.59 | ND | 57.52 ± 2.37 | 60.90 ± 1.51 |
| Fruc + C2 | 1.64 ± 0.20 | 14.33 ± 0.42 | 10.20 ± 2.41 | -14.26 ± 8.33 | ND | 38.66 ± 7.15 | ND | 40.27 ± 15.41 | 44.08 ± 5.90 |
| Fruc + C3 | 1.06 ± 0.06 | NM | 5.01 ± 2.76 | 15.60 ± 0.52 | -57.53 ± 2.03 | 6.67 ± 0.59 | 76.51 ± 0.40 | 2.99 ± 0.64 | 2.79 ± 0.52 |
| Fruc + C4 | 1.69 ± 0.06 | 27.56 ± 2.80 | 6.25 ± 1.05 | 19.31 ± 0.54 | ND | -55.13 ± 1.15 | ND | 125.49 ±1.89 | 83.08 ± 0.44 |
| Fruc + C5 | 1.08 ± 0.11 | NM | 46.15 ± 3.15 | 8.78 ± 1.15 | ND | 7.04 ± 1.05 | -10.95 ± 0.78 | 44.31 ± 5.3 | 41.43 ± 3.33 |
| Fruc + C6 | 0.17 ± 0.1 | NM | 4.02 ± 1.26 | ND | ND | ND | ND | -1.96 ± 0.54 | NP |

Positive values represent production and negative values represent consumption. Error represents one standard deviation among triplicates. ND: not detected; NM: not measured; NP: not produced. The pH value of the test was 5.5 ± 0.02. Fruc: fructose; C2: acetate; C3: propionate; C4: *n*-butyrate; C5: *n*-valerate; C6: *n*-caproate.

**Table S2** Maximum $OD_{600}$ values, final concentration of products, and specificities of lactate and *n*-caproate in cultures of strain 7D4C2 grown at different pH values and temperatures.

| pH / Temp ( / °C) | Max $OD_{600}$ | Growth rate* ($d^{-1}$) | $H_2$ production (mmol $L^{-1}$) | Lactate concentration (mmol C $L^{-1}$) | Acetate concentration (mmol C $L^{-1}$) | *n*-Butyrate concentration (mmol C $L^{-1}$) | *n*-Caproate concentration (mmol C $L^{-1}$) | Lactate carboxylate specificity (%) | *n*-Caproate carboxylate specificity (%) | Average fructose consumption (mmol C $L^{-1}$ $d^{-1}$) |
|---|---|---|---|---|---|---|---|---|---|---|
| 4.5 / 30 | 0.43 | NA | 4.40 | 0.00 0.00 | 8.15 10.6 | -57.02 -51.18 | 93.18 108.24 | 0.00 0.00 | 91.08 92.78 | -1.9 -0.2 |
| 5.0 / 30 | 1.25 1.29 | 0.63 | 10.74 11.29 | 2.01 3.753 | 9.66 10.09 | -81.96 -78.6 | 138.02 142.36 | 1.30 2.47 | 90.88 92.42 | -7.32 -7.03 |
| 5.2 / 30 | 1.15 1.69 | 0.50 | 23.22 24.29 | 4.58 5.00 | 12.49 14.33 | -78.55 -78.41 | 129.16 146.66 | 3.01 3.13 | 88.32 88.36 | -17.34 -25.39 |
| 5.5 / 30 | 1.56 1.58 | 0.34 | 24.28 24.89 | 9.23 11.91 | 17.14 17.56 | -70.24 -68.86 | 137.04 139.69 | 5.55 6.36 | 82.51 83.91 | -24.91 -24.91 |
| 6.0 / 30 | 1.64 1.98 | 1.3 | 18.09 20.32 | 60.52 61.16 | 17.94 18.38 | -75.03 -65.75 | 119.81 123.03 | 30.26 30.45 | 60.30 60.87 | -37.04 -37.04 |
| 6.5 / 30 | 1.03 1.23 | 0.42 | 12.58 15.07 | 84.62 90.39 | 21.44 32.90 | -44.41 -38.44 | 99.24 135.72 | 33.41 42.82 | 47.02 53.59 | -24.45 -24.45 |
| 7.0 / 30 | 0.85 1.35 | 0.69 | 7.98 8.11 | 75.05 102.04 | 14.81 25.29 | -43.37 -35.43 | 54.02 75.88 | 50.22 52.16 | 37.34 37.54 | -24.45 -24.45 |
| 7.5 / 30 | 0.90 0.93 | 0.48 | 9.41 11.67 | 93.50 97.03 | 11.17 18.90 | -36.55 -36.37 | 62.83 68.45 | 52.62 55.82 | 37.12 37.51 | -24.45 -24.45 |
| 8.0 / 30 | 1.00 1.31 | 0.98 | 5.94 8.01 | 87.51 103.54 | 13.59 16.69 | -29.86 -27.34 | 37.28 49.67 | 60.94 63.24 | 26.94 29.23 | -24.45 -24.45 |
| 9.0 / 30 | 1.02 1.05 | 0.85 | 7.94 10.63 | 97.12 103.01 | 17.34 19.60 | -19.48 -16.70 | 35.64 37.77 | 64.23 64.70 | 23.55 23.75 | -16.30 -16.30 |
| 6.0 / 22.5 | 1.66 1.77 | 0.45 | 14.72 18.97 | 35.84 39.52 | 16.07 17.08 | -60.12 -54.62 | 113.57 115.01 | 21.61 23.17 | 67.41 68.57 | -10.14 -9.78 |
| 6.0 / 27 | 1.59 1.61 | 0.88 | 18.42 19.52 | 52.43 57.52 | 15.25 15.55 | -54.14 -52.54 | 96.32 99.62 | 31.34 33.95 | 56.86 59.55 | -27.29 -27.29 |
| 6.0 / 30 | 1.37 1.39 | 0.95 | 17.66 17.94 | 38.03 38.07 | 15.87 15.95 | -56.75 -56.72 | 105.83 108.94 | 23.36 23.81 | 66.26 66.85 | -27.29 -27.29 |
| 6.0 / 37 | 1.35 1.36 | 0.96 | 22.41 33.94 | 21.44 26.42 | 26.13 26.25 | -43.84 -40.97 | 88.53 88.54 | 15.75 18.71 | 62.70 65.05 | -45.48 -0.45.48 |
| 6.0 / 42 | 1.40 1.40 | 0.87 | 19.87 22.90 | 50.39 50.93 | 19.20 20.11 | -39.41 -36.38 | 69.44 79.18 | 33.90 36.24 | 49.95 52.71 | -45.48 -0.45.48 |

Negative values denote consumption. NA: not available

*Growth rate given for the exponential phase (**see Figure S3**), calculated by plotting Ln($OD_{600}$) *vs*. time.

**Table S3** Average nucleotide identity (ANI) and alignment fraction (AF) values between strain 7D4C2 and the most similar type strains of the order Clostridiales (heterotypic synonym of Eubacteriales) and unclassified species which are closely related; as calculated by JSpeciesWS.

| **Genome** | **ANI** | **AF** |
|---|---|---|
| *Clostridium* sp. W14A | 97.64 | 82.28 |
| *Caproicibacter fermentans* EA1 | 97.34 | 81.20 |
| *Caproiciproducens* sp. NJN-50 | 78.52 | 44.36 |
| *Clostridium* sp. KNHs216 | 71.14 | 32.50 |
| *Ethanoligenens harbinense* YUAN-3 | 69.65 | 15.30 |
| *Caproiciproducens galactitolivorans* BS-1 | 69.34 | 28.22 |
| *Anaeromassilibacillus senegalensis* mt9 | 69.21 | 21.65 |
| *Oscillibacter valericigenes* Sjm18-20 | 68.74 | 16.15 |
| *Clostridium jeddahense* JCD | 68.30 | 22.78 |
| *[Clostridium sporosphaeroides* VPI 4527 | 68.20 | 23.71 |
| *[Clostridium] leptum* VPI T7-24-1 | 67.55 | 15.78 |
| *Clostridium minihomine* Marseille-P4642 | 67.40 | 19.62 |
| *Clostridium merdae* Marseille-P2953 | 66.78 | 19.09 |
| *Clostridium kluyveri* K1 | 65.32 | 6.05 |
| *Christensenella minuta* DSM 22607 | 64.67 | 8.00 |
| *Eubacterium limosum* SA11 | 64.11 | 8.54 |

**Table S4** Comparison of carbohydrates oxidized by strain 7D4C2, *C. galactitolivorans*, and *C. fermentans*.

| Substrate | strain 7D4C2 | *C. galactitolivorans* | *C. fermentans* |
|---|---|---|---|
| N-Acetyl-D- Glucosamine | + | - | |
| Adonitol | - | - | |
| Amygdalin | - | - | |
| D-Arabitol | + | - | |
| Arbutin | - | - | |
| D-Cellobiose | + | + | + |
| Dextrin | + | + | |
| Dulcitol | - | + | |
| i-Erythritol | - | - | |
| D-Fructose | + | + | + |
| L-Fucose | - | + | |
| D-Galactose | - | + | + |
| Gentiobiose | + | - | |
| D-Gluconic Acid | - | - | |
| α-D-Glucose | + | + | |
| Glycerol | + | + | - |
| m-Inositol | - | + | |
| α-D-Lactose | - | - | |
| Maltose | + | - | |
| D-Mannitol | + | + | + |
| D-Mannose | + | + | + |
| D-Melezitose | + | - | |
| D-Melibiose | + | - | |
| D-Raffinose | - | - | |
| L-Rhamnose | - | - | |
| Salicin | - | - | |
| D-Sorbitol | + | + | + |
| Sucrose | + | - | |
| D-Trehalose | + | - | |
| Turanose | + | - | |

**"+"** (highlighted in green) indicate that a substrate is metabolized and **"-"** indicate that a substrate is not metabolized, according to the AN MicroPlate$^{TM}$ from Biolog (Hayward, CA) for strain 7D4C2 (**Figure S7**), and the data reported for *C. galactitolivorans* in Kim *et al*., 2015 and for *C. fermentans* in Flaiz *et al*. Blank: not reported.

**Table S5** Composition of the supplemented basal medium, the modified Wolfe's trace minerals solution, and the vitamin solution. For the serial dilutions in soft agar, the medium contained the substrates as summarized in **Figure S1**.

| **1. Supplemented Basal medium (1 L)** | | **2. Modified Wolfe's Trace minerals (1 L)** | | **3. Vitamin solution 2x (1 L)** | |
|---|---|---|---|---|---|
| NaCl | 1.1 g | EDTA | 500 mg | Pyridoxine hydrochloride | 20 mg |
| $NH_4Cl$ | 0.474 g | $MgCl_2 \cdot 6H_2O$ | 3.63 g | Riboflavin | 10 mg |
| $CaCl_2 \cdot 2H_2O$ | 0.1 g | $MnCl_2 \cdot 4H_2O$ | 427.1 mg | Nicotinic acid | 10 mg |
| $MgCl_2 \cdot 6H_2O$ | 0.065 g | NaCl | 1 g | Folic acid | 4 mg |
| $KH_2PO_4$ | 0.15 g | $CoCl_2 \cdot 6H_2O$ | 180 mg | Calcium pantothenate | 10 mg |
| $Na_2CO_3$ | 0.032 g | $CaCl_2$ (anhydrous) | 100 mg | P-amino benzoic acid | 10 mg |
| Trace mineral Wolfe modified | 20 mL | $ZnCl_2$ | 210.9 mg | D-biotin | 4 mg |
| Na-Butyrate | 3.3 g | $CuCl_2 \cdot 2H_2O$ | 14.64 mg | Thioctic acid | 10 mg |
| DI $H_2O$ | 856 mL | $H_3BO_3$ | 10 mg | Vitamin B12 | 10 mg |
| **Sparge 30-40 min $N_2$:$CO_2$ (80:20)** | | $Na_2MoO_4 \cdot 2H_2O$ | 10 mg | Thiamine HCl | 10 mg |
| L-cysteine HCl | 0.47 g | $Na_2SeO_3$ (anhydrous) | 1 mg | 2-mercaptoethanesulfonic acid | 4 mg |
| $FeCl_2$ (10g/L stock solution) | 0.125 mL | $Na_2WO_4 \cdot 2H_2O$ | 10 mg | | |
| **Autoclave 121ºC, 20min** | | $NiCl_2 \cdot 6H_2O$ | 20 mg | | |
| Yeast extract (10% w/v)[a] | 20 mL | | | | |
| Fructose or glucose (250 g/L)[a] | 22 mL | | | | |
| Vitamin solution 2x[a] | 2 mL | | | | |
| MES 1M[a] | 100 mL | | | | |
| **Adjust to pH 5.5-5.6** | | | | | |

[a]filter sterilized
